# The tubulin poly-glutamylase complex, TPGC, is required for phosphatidyl inositol homeostasis and cilium assembly and maintenance

**DOI:** 10.1101/2025.03.03.641315

**Authors:** Binshad Badarudeen, Hung-Ju Chiang, Loren Collado, Lei Wang, Irma Sanchez, Brian David Dynlacht

## Abstract

The tubulin poly-glutamylase complex (TPGC) is comprised of TTLL1 and at least five associated proteins that promote the addition of glutamate residues to tubulin tails of microtubules. Despite its discovery two decades ago, the enzyme has been refractory to characterization owing to its complex multimeric nature and the inability to detect poly-glutamylase activity after assembling the six-subunit complex. We now show that TPGC is the key enzyme driving centriolar and ciliary poly-glutamylation. We identified two novel TPGC subunits, TBC1D19 and KIAA1841, and showed that both components play an essential role in the assembly of the eight-subunit holo-enzyme. Remarkably, we were able to reconstitute the activity of TPGC with all eight subunits. TBC1D19 and KIAA1841 were essential for assembly and activity, and loss of TBC1D19 strongly compromised multiple tubulin modifications, including axonemal poly-glutamylation. TBC1D19 loss abolished transport of Arl13b and other ciliary membrane proteins, abrogating primary cilium assembly. Structural modeling revealed an essential role for TBC1D19 and KIAA1841 in complex assembly, microtubule binding, and preferential poly-glutamylation of α-tubulin. We found that TBC1D19 loss abrogated the ciliary localization of phosphatidyl inositol phosphatase, INPP5E, triggering cilium instability. Ciliogenesis in TBC1D19 null cells could be restored through inhibition of a specific phosphatidyl inositol phosphate (PIP) kinase, PIP5K1c, suggesting that TBC1D19 is required to instigate and maintain PIP homeostasis during ciliogenesis. Collectively, our data show that TPGC is a multi-functional enzyme essential for cilium assembly and maintenance.

## Introduction

Recent studies have suggested that tubulin poly-glutamylation regulates a panoply of cellular processes. For example, poly-glutamylation modulates microtubule (MT) dynamics by stimulating severing enzymes (Sharma, Bryant et al. 2007, Lacroix, van Dijk et al. 2010, Valenstein and Roll-Mecak 2016, Szczesna, Zehr et al. 2022) or by altering recruitment of kinesins, dynein, or <u>m</u>icrotubule <u>a</u>ssociated <u>p</u>roteins (MAPs), which can promote MT assembly (Bonnet, Boucher et al. 2001, Janke and Bulinski 2011, Sirajuddin, Rice et al. 2014). Interestingly, *in vitro* studies suggest that poly-glutamylation has a rheostat-like role in modulating microtubule cleavage by spastin, since precise levels of glutamylation are required for optimal enzyme activity (Valenstein and Roll-Mecak 2016).

Tubulin glutamylation also plays a role in the assembly of flagella and primary cilia. Primary cilia are antenna-like, microtubule-based projections found on the surface of most eukaryotic cells. In the early stages of cilium assembly, a protein (CP110) found at the distal end of centrioles is destroyed at the mother centriole, triggering microtubule growth from the basal body and generating a microtubular axoneme that supports the cilium and promotes intra- flagellar transport (IFT) required for its assembly (Spektor, Tsang et al. 2007, Sanchez and Dynlacht 2016). The nascent cilium then migrates apically and docks with the plasma membrane. As cilia assemble, axonemal microtubules (MTs) are de-tyrosinated and decorated with several post-translational modifications (PTMs), including acetylation and glutamylation, and these modifications are thought to provide the underlying stability of the axoneme. Conversely, MTs are de-acetylated during disassembly. However, the functional and mechanistic links between microtubule glutamylation and axoneme assembly and dynamics in mammalian cells are not completely understood, and functional roles in both cilium assembly and disassembly have been ascribed to glutamylation, suggesting complex and, potentially, context-dependent roles for this modification (Lee, Silhavy et al. 2012, Hong, Wang et al. 2018, Ki, Kim et al. 2020, Latour, Van De Weghe et al. 2020).

Microtubule glutamylation is catalyzed by the tubulin tyrosine ligase-like (TTLL) family of enzymes comprised of 13 members. Several family members (TTLL1, TTLL5, TTLL6, and TTLL9) have been shown to promote ciliary axonemal poly-glutamylation (Wloga, Joachimiak et al. 2017). However, understanding the biological impact of microtubule (MT) poly- glutamylation has been challenging, in part due to redundancy within the TTLL family and species- and tissue-specific roles for glutamylation. As examples, depletion of TTLL1 in mice leads to shortened sperm flagella but normal respiratory cilia (Janke, Rogowski et al. 2005, Ikegami, Sato et al. 2010, Vogel, Hansen et al. 2010). Moreover, depletion of another family member, TTLL5, in human RPE1 cells or mice provoked cilium disassembly and sperm motility defects, respectively (Lee, He et al. 2013, He, Ma et al. 2018).

In contrast with all other glutamylases, TTLL1 is not active as a monomer and lacks a MT-binding domain universally found in other family members (van Dijk, Rogowski et al. 2007, Garnham, Vemu et al. 2015), suggesting that one or more associated proteins modulate its function by directing its sub-cellular localization, by recruiting the holo-enzyme to MTs, or by allosterically activating the enzyme. To that end, biochemical purification of a mammalian tubulin poly-glutamylation complex (TPGC) two decades ago identified TTLL1 and four additional associated proteins (LRRC49/CSTPP2, NICN1, TPGS1, and TPGS2) (Janke, Rogowski et al. 2005). More recent studies revealed a sixth TPGC subunit, C11orf49/CSTPP1, and ablation experiments have confirmed the importance of TTLL1-associated proteins (Wang, Paudyal et al. 2022). For example, loss of C11orf49/CSTPP1 or LRRC49/CSTPP2 compromised tubulin poly-glutamylation, resulting in aberrant nuclear envelope assembly and nuclear lobulation, and null cells exhibited abnormal persistence of primary cilia in growing cells (Wang, Paudyal et al. 2022). Moreover, *TPGS1^-/-^* mice exhibited selective loss of α-tubulin poly-glutamylation and defects in transport of synaptic vesicles, which are kinesin cargoes, resulting in impaired neuronal synaptic transmission (Ikegami, Heier et al. 2007).

The observation that TTLL1 stably integrates into a large multimeric complex suggests an intricate role for TPGC in precisely regulating and fine-tuning microtubule poly-glutamylation and perhaps other cellular processes, and given the pervasive links between genetic defects associated with aberrant tubulin modifications and human disease, it is essential to understand the underlying mechanisms. In an effort to comprehensively define all aspects of the TPGC, we affinity-purified this complex and identified two novel subunits, TBC1D19 and KIAA1814, required to assemble and activate the holo-enzyme. We observed that TBC1D19 was required for tubulin poly-glutamylation as well as cilium assembly and maintenance, in part by regulating entry of Arl13b and INPP5E into the primary cilium, promoting its stability. Structural modelling of the TPGC holo-enzyme confirmed our findings and provided novel insights into why TTLL1 activity requires both of these additional factors for robust microtubule-binding, positioning of the carboxy-terminal tail of α-tubulin within its catalytic cleft, and poly-glutamylase activity. Our findings provide important new insights into mechanisms regulating the biological activity of TPGC, a major cellular tubulin poly-glutamylase.

## Results and Discussion

### Identification of TBC1D19 and KIAA1841 as integral components of the TPGC holo-enzyme

Our previous studies revealed that ablation of two proteins, C11orf49/CSTPP1 or LRRC49/CSTPP2, significantly reduced overall levels of microtubule poly-glutamylation, but neither protein proved essential for this activity (Wang, Paudyal et al. 2022). However, when we attempted to assemble the entire complex through simultaneous expression of all six TPGC subunits in mammalian cells, we were unable to reconstitute poly-glutamylase activity. These findings prompted us to search for additional TPGC-associated factors required to assemble and activate the enzyme. The BioGRID repository revealed two candidates (TBC1D19 and KIAA1841/SANBR), which had been identified in proteome-wide affinity capture screens with multiple TPGC subunits (Huttlin, Bruckner et al. 2021), although these proteins remain uncharacterized (TBC1D19) or partially characterized (KIAA1841/SANBR; (Zheng, Matthews et al. 2021). We generated a stable human RPE1 cell line expressing GFP-TBC1D19 protein (Fig. 3a, d), purified proteins associated with the fusion protein through GFP-trap chromatography, and performed mass spectrometric sequencing, which captured all known subunits of TPGC, except for TPGS2 (Fig. 1a). We also confirmed these interactions through co-immunoprecipitation with GFP-TBC1D19 and found that KIAA1841/SANBR was indeed a stable member of this complex (Figs. 1b and S1a). KIAA1841/SANBR, a protein implicated in class switch recombination in B cells (Zheng, Matthews et al. 2021), was previously identified in high-throughput affinity-capture studies as an interacting partner of multiple TPGC components, including TTLL1, NICN1, C11orf49/CSTPP1, LRRC49/CSTPP2, and TPGS1 (Huttlin, Bruckner et al. 2021). These studies suggested that TBC1D19 and KIAA1841/SANBR are *bona fide* components of the TPGC complex. Since we have not reproducibly identified additional interacting partners in multiple GFP-trap affinity purification experiments, we propose that the TPGC consists of eight subunits and term this complex, comprised of C11orf49/CSTPP1, LRRC49/CSTPP2, TTLL1, TPGS1, TPGS2, TBC1D19, KIAA1841, and NICN1, the TPGC holo-enzyme or holo-TPGC.

**Figure 1.**
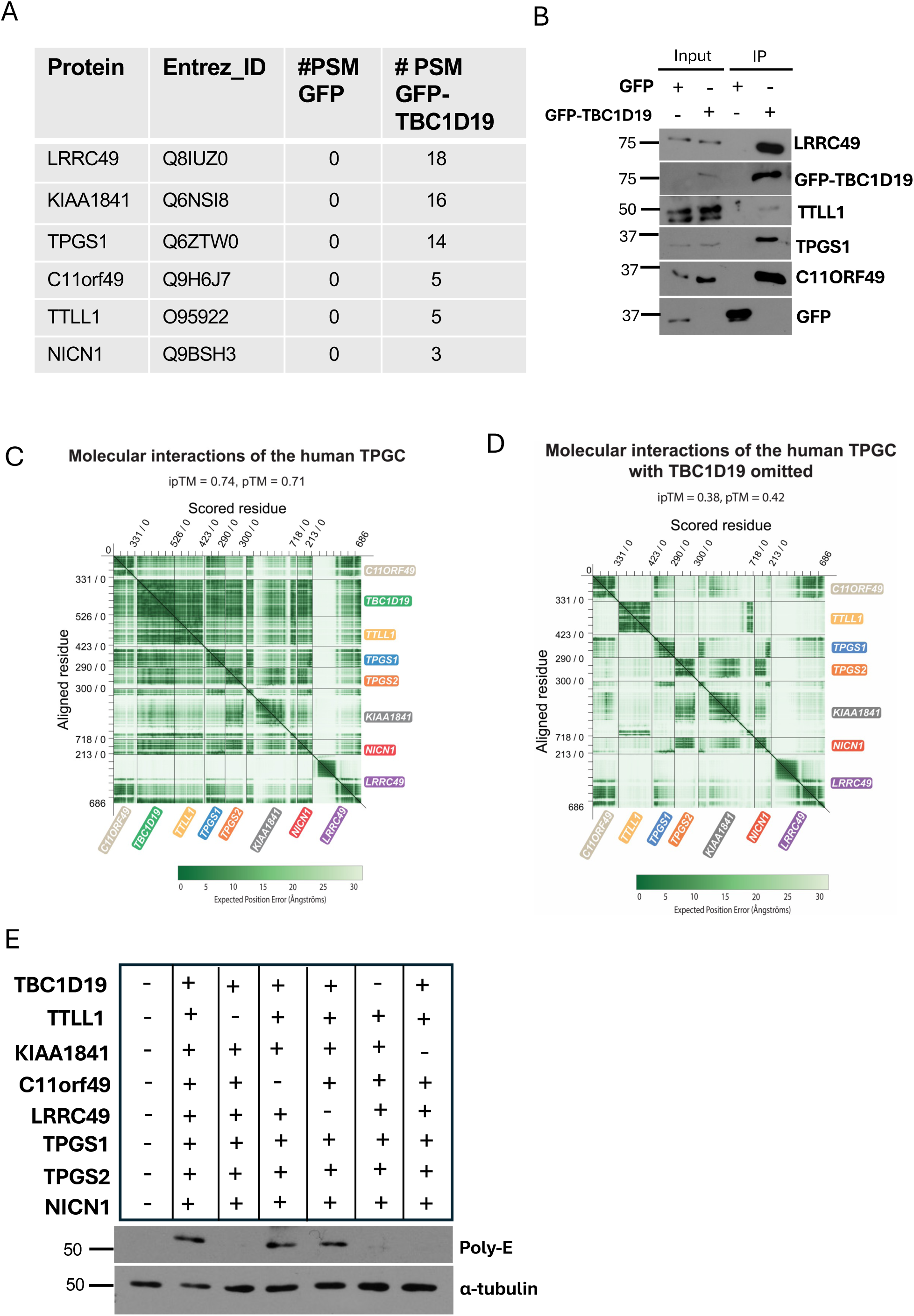
TBC1D19 regulates the assembly of tubulin poly-glutamylase complex. **A**) Table showing the number of peptides observed after GFP-trap purification of RPE-1 cells expressing GFP only (control) or GFP-TBC1D19 and mass spectrometric sequencing. **B**) RPE-1 cells stably expressing GFP or GFP-TBC1D19 were subjected to immunoprecipitation with GFP-trap beads. Input and immunoprecipitates were immunoblotted with the indicated antibodies. **C**) Predicted aligned error (PAE) matrix of human TPGC obtained with AlphaFold3. **D**) PAE matrix of human TPGC after omission of TBC1D19. **E**) Robust poly-glutamylation requires holo-TPGC. Extracts of 293T cells programmed to express combinations of epitope-tagged TBC1D19, KIAA1841, TTLL1, NICN1, TPGS2, TPGS1, C11ORF49, and LRRC49 (indicated by “+”), were immunoblotted with antibodies as shown.

### Modeling the TPGC holo-enzyme reveals a core and regulatory “arm”

We used AlphaFold3 to model high-confidence inter-molecular interactions within holo-TPGC and TPGC sub-complexes (Abramson, Adler et al. 2024). Several lines of evidence suggest that this approach is ideally suited for accurately modeling this complex. First, we showed that our methods were able to model tubulin dimers, which could be iteratively added to form tubulin protomers (Fig. S1). Second, cryo-EM studies have elucidated the structures of several TTLL enzymes bound to microtubules, including TTLL7, which docks to the body of α-tubulin through its microtubule-binding domain (MBD), allowing the enzyme to preferentially bind and glutamylate the carboxy-terminal tail of β-tubulin (Garnham, Vemu et al. 2015). AlphaFold3 modeling successfully captured selective interactions between the catalytic domain of TTLL7 and the β-tubulin tail, as well as interactions between the microtubule-binding domain and α-tubulin (Fig. S1c). These examples (and others; see below) provided confidence in our ability to accurately model TTLL proteins and associated tubulin dimers, and they prompted us to use AlphaFold3 to make key structural predictions regarding holo-TPGC (Fig. 1c). Remarkably, when we modeled the intra- and inter-protein interactions within holo-TPGC, we were able to predict high-confidence structures able to accommodate all eight subunits of the complex (Fig. 2a). Modeling led to another key prediction: when we attempted to assemble the holo-enzyme without TBC1D19, nearly all high-confidence inter-protein interactions were nullified (see below), suggesting that the protein is an essential assembly factor (Fig. 1d).

**Figure 2.**
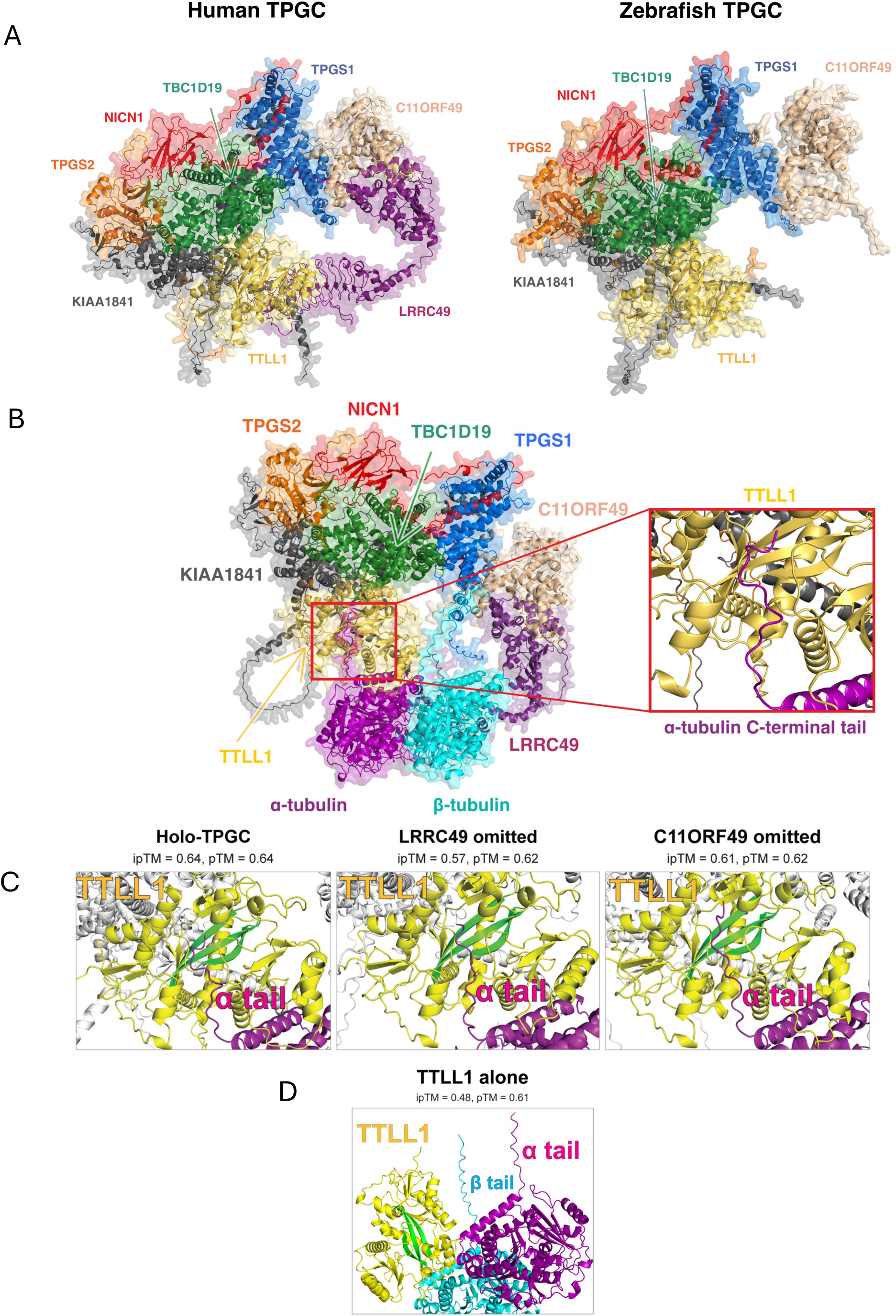
Structural modeling of holo-TPGC complex bound to tubulin. **A**) 3D structures of Human and Zebrafish TPGC complexes were predicted with AlphaFold3 and visualized with PyMOL. **B**) 3D structure of human TPGC complex with α-tubulin and β-tubulin. Image inset shows the positioning of α-tubulin carboxy-terminal tail (magenta) within the catalytic cleft of TTLL1. **C**) Enlarged view of 3D structures from panel B showing the insertion of α-tubulin tail within the catalytic cleft of TTLL1 assembled into holo-TPGC or after omission of C11ORF49 or LRRC49 from the holo-enzyme. Active site β sheets in TTLL1 conserved with TTLL4, 5, 6, and 7 are indicated (green). **D)** Model showing TTLL1 alone with tubulin protomer.

TTLL1 is not active as a monomer and does not have a recognizable microtubule-binding domain (van Dijk, Rogowski et al. 2007, Garnham, Vemu et al. 2015). Further, TTLL family members preferentially poly-glutamylate microtubules rather than tubulin monomers (van Dijk, Rogowski et al. 2007). To explore the structural bases for these observations, we used AlphaFold3 to model interactions between TTLL1 or holo-TPGC and tubulin dimers. Modeling TTLL1 alone with tubulin dimers failed to produce high-confidence interactions, and indeed, all such predictions indicated that the a-tubulin carboxy-terminal tail was located far from the active site of TTLL1 (Fig. 2d). In contrast, when we docked holo-TPGC onto tubulin dimers, TTLL1 form extensive contacts with α-tubulin, but not β-tubulin, consistent with the observation that the glutamylase preferentially modifies the α-tubulin carboxy-terminal tail. Although tubulin carboxy-terminal tails are thought to be intrinsically disordered (Natarajan, Gadadhar et al. 2017), we nevertheless obtained high-confidence models demonstrating that the a-tubulin tail can dock within the catalytic cleft of TTLL1 when assembled into holo-TPGC (Fig. 2b, Video 2). The carboxy-terminal tail of ý-tubulin was unable to engage the active site cleft of TTLL1, further suggesting a mechanism whereby the enzyme preferentially modifies a-tubulin. Since TTLL1 alone was unable to dock with the tubulin dimer, our models suggest that assembly of holo-TPGC is required to position the a-tubulin tail within the active site of TTLL1.

Importantly, our structural predictions with TTLL1 recapitulated the crystallographic data for TTLL4, as well as homology models generated for TTLL5, TTLL6, and TTLL7, with respect to the conserved active site and β-sheet motifs near the catalytic cleft of the enzyme (Fig. S1d; (Natarajan, Gadadhar et al. 2017). Notably, although each of these family members shared homology within their active sites, we were unable to obtain high-confidence structural models of holo-TPGC when we substituted TTLL6 for TTLL1, indicating that residues in TTLL1 outside the catalytic cleft were essential for its interactions with other subunits within holo-TPGC. These observations further attest to the ability of AlphaFold3 to model holo-TPGC.

Next, we investigated a role, if any, for subunits within holo-TPG as regulators of the interaction between TTLL1 and the C-terminal tail of α-tubulin. Further, to gain insights into the role of individual proteins within holo-TPGC when docked to a tubulin protomer, we predicted the impact of removing one subunit at a time using a “leave-one-out” strategy. Modeling showed that TBC1D19 formed extensive interactions with TTLL1, and the docking of holo-TPGC and tubulin dimers absolutely required TBC1D19, TTLL1, and KIAA1841 (Figs. 1c and S1e). Remarkably, omission of either KIAA1841 or TBC1D19 produced models wherein the α-tubulin carboxy-terminal tail was unable to dock with TTLL1, and indeed, the tail was again predicted to be located at a substantial distance from TTLL1 (Fig. S1e). Our data therefore suggest that both TBC1D19 and KIAA1841 are required for TTLL1 to properly dock to α-tubulin. In striking contrast, omission of either LRRC49/CSTPP2 or C11ORF49/CSTPP1 did not abrogate TTLL1 docking to the α-tubulin tail, indicating that these proteins may not be essential for promoting interactions between TTLL1 and α-tubulin (Fig. 2c). Instead, these proteins may be required to fine-tune the poly-glutamylation activity of holo-TPGC, consistent with our data comparing poly-E levels within TBC1D19, TTLL1, and C11orf49 KO cell lines (see below; Fig. 5a and b).

Further, we used AlphaFold3 to assess the contributions of each subunit to the integrity of holo-TPGC. Our modeling predicted that loss of TBC1D19 would lead to a catastrophic collapse of holo-TPGC and abolish the docking of the α-tubulin tail to TTLL1. In sharp contrast with omission of TBC1D19 or KIAA1841, our modeling suggested that C11orf49/CSTPP1 and LRRC49/CSTPP2 interacted with one another to form a distinct module or “arm,” permitting it to assemble independently from a “core” complex composed of TBC1D19, TTLL1, TPGS1/2, NICN1, and KIAA1841. Importantly, this modeling agrees with analysis of C11orf49/CSTPP1 and LRRC49/CSTPP2 knock-out cells, wherein ablation of C11orf49/CSTPP1 or LRRC49/CSTPP2 had a reciprocal and dramatic impact on the stability of each other but had a muted effect on TPGS1 and TPGS2 (Wang, Paudyal et al. 2022). Interestingly, loss of C11orf49/CSTPP1 or LRRC49/CSTPP2 led to decreased stability of TTLL1, potentially explaining the diminution of poly-glutamylated microtubules in these knockout lines (Wang, Paudyal et al. 2022). Further, loss of either LRRC49/CSTPP2 or C11orf49/CSTPP1 moderately altered the positioning of the catalytic cleft with respect to the α-tubulin tail (Fig. 2c), but their omission did not displace the tail from the active site, in contrast with the loss of TBC1D19, suggesting a plausible explanation for the degree to which poly-glutamylation is affected in the respective knockouts (Fig. 5a and b; (Wang, Paudyal et al. 2022).

TPGC components are well-conserved in vertebrates, including zebrafish, allowing us to model the orthologous complex. Although a zebrafish LRRC49 ortholog has been previously nominated, this protein is a fraction of the size of the human protein (197 versus 686 amino acids), and all homologies are restricted to short stretches within the leucine-rich repeat (LRR), a widely occurring motif that frequently adopts a horseshoe configuration (Kobe and Deisenhofer 1994). Moreover, addition of the putative zebrafish ortholog to our models of holo-TPGC failed to yield high-confidence inter-molecular interactions in the predicted aligned error (PAE) matrix. We therefore rejected the zebrafish protein as an ortholog and omitted it from all subsequent models of the zebrafish complex. Interestingly, despite the lack of a zebrafish LRRC49/CSTPP2 ortholog and 61% amino acid identity overall (when averaged across all seven remaining subunits), intra- and inter-molecular interaction maps for human and zebrafish holo-TPGC strongly resembled one another, suggesting both structural and functional conservation (Fig. 2a). In both mammalian and zebrafish holo-TPGC, the TTLL1 catalytic cleft was positioned near E445 of the α-tubulin carboxy-terminal tail, an acceptor for the branch-point glutamate and subsequent addition of poly-E chains. As before, omission of zebrafish TBC1D19 abolished all inter-subunit interactions within the complex. Thus, our key observations regarding human holo-TPGC were mirrored by modeling the zebrafish complex, further strengthening our conclusions regarding complex architecture (Fig. 2a).

Next, we asked whether we could reconstitute the TPGC complex and induce glutamylation *in vitro*. Prior efforts to assemble an active enzyme with six subunits (C11orf49/CSTPP1, LRRC49/CSTPP2, TTLL1, TPGS1, TPGS2, and NICN1) were unsuccessful. We therefore expressed all eight subunits in 293T cells and analyzed cellular glutamylation. Remarkably, we were able to reconstitute robust TPG glutamylation activity, for the first time, through simultaneous expression of all eight subunits, including TBC1D19 and KIAA1841 (Fig. 1e). Omission of C11ORF49/CSTPP1 or LRRC49/CSTPP2 had little or no impact on activity in this system (Fig. 1e). Importantly, we observed a striking diminution of glutamylation activity when we omitted either TBC1D19 or KIAA1841. Moreover, TTLL1, TPGS2 and NICN1 were destabilized when all subunits except TBC1D19 were expressed (Fig. S1b). We also observed that TPGS1 was lost when C11ORF49 was omitted. Overall, our TPGC reconstitution experiment confirmed our structural modeling approach and verified the requirement for all eight components to functionally assemble TPGC and induce robust glutamylation activity.

In summary, our modeling indicates that assembly of holo-TPGC is needed to stabilize interactions with tubulin dimers and ensure proximity between critical active site residues in TTLL1 and the α-tubulin carboxy-terminal tail to promote catalysis, explaining why TTLL1 alone does not independently bind or glutamylate MTs. Our modeling further suggests that (1) the TPGC “core” can assemble independently from the module consisting of C11orf49/CSTPP1 and LRRC49/CSTPP2; (2) TBC1D19 lies at the center of the TPGC core and is essential for assembly and stabilization of holo-TPGC, and loss of TBC1D19 abolishes complex assembly; (3) C11orf49/CSTPP1 acts as a linker to connect LRRC49/CSTPP2 with the “core”; (4) KIAA1841 is required to dock TPGC onto tubulin dimers; and (5) assembly of holo-TPGC is required to ensure proper orientation and positioning of critical catalytic residues of TTLL1 in proximity to branch-point glutamate acceptors within the α-tubulin tail.

### TBC1D19 is required for cilium assembly and stability

Given the pivotal role of TBC1D19 as an essential assembly factor, we focused all subsequent studies on elucidating its function. First, we examined its localization using our stable GFP-TBC1D19 RPE1 cell line and found that the protein was enriched at centriolar satellites, electron-dense structures surrounding the centrosome, consistent with our studies of C11orf49/CSTPP1 and LRRC49/CSTPP2 (Wang, Paudyal et al. 2022)(Fig. 3a). Further, when cells were induced to ciliate, TBC1D19 localized at a region proximal to the basal body (Fig. 3b). The localization of TBC1D19 indicated a possible role in primary cilium assembly. To elucidate the role of TBC1D19 in poly-glutamylation and ciliogenesis, we used CRISPR to knock out the gene in human RPE1 cells. Since our modeling predicted that TBC1D19 would have an impact on TTLL1 function, we also ablated *TTLL1* and compared the resulting phenotypes in both knockouts with *C11orf49/CSTPP1* and *LRRC49/CSTPP2* null cells (Fig. S2a and b; (Wang, Paudyal et al. 2022). Consistent with our modeling studies, we found that ablation of TBC1D19 resulted in the near-disappearance of multiple TGPC subunits and reductions in tubulin poly-glutamylation in growing and serum-starved cells (Fig. 3c). We were able to restore poly-glutamylation and native levels of expression for each of these proteins upon expression of TBC1D19 in knock-out cells, confirming that their loss resulted specifically from ablation of this protein (Fig. 3d). Proper localization of C11orf49/CSTPP1 was not dependent upon TBC1D19, and likewise, TBC1D19 was not impacted by loss of TTLL1 (Fig. S2c), consistent with a model in which TPGC subunits can independently localize to centriolar satellites, most likely through an association with PCM1, which could serve as a platform to stabilize or promote complex assembly (Wang, Paudyal et al. 2022).

**Figure 3.**
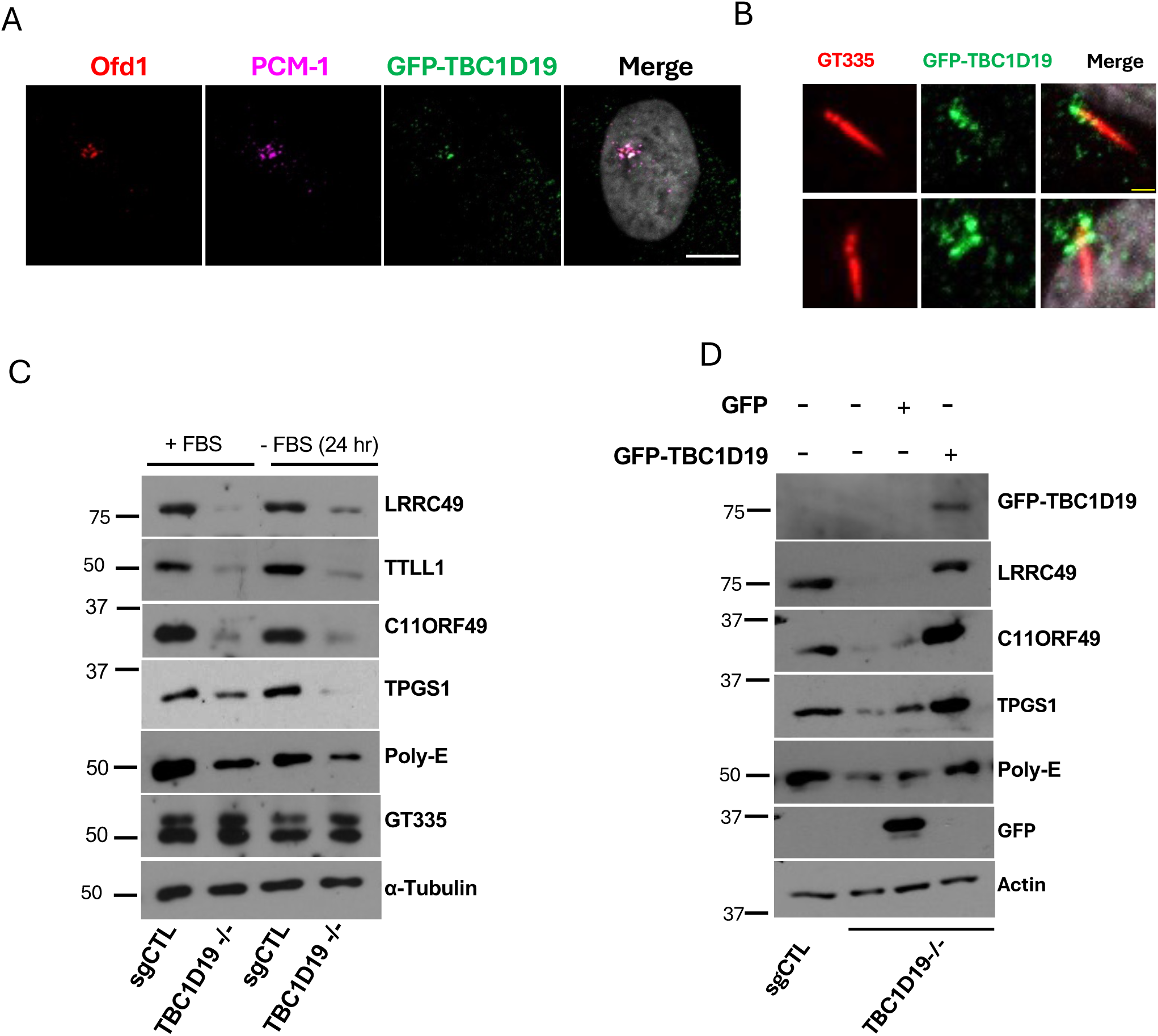
TBC1D19 regulates cellular poly-glutamylation. **A**) Growing RPE-1 cells stably expressing GFP-TBC1D19 were fixed and immuno-stained with antibodies against PCM-1 and Ofd1. DAPI was used to stain nuclei. Scale bar, 10 µm. **B**) RPE-1 cells stably expressing GFP-TBC1D19 were serum starved for 24 hr, fixed, and stained with antibodies against GT335 and GFP. Scale bar, 1µm. **C**) sgRNA control and TBC1D19 knockout cells were subjected to western blot analysis with the indicated antibodies. **D**) TBC1D19 knockout cells were infected with GFP control or GFP-TBC1D19 vector for 72hr and subjected to western blot analysis with the indicated antibodies.

Interestingly, in contrast with LRRC49/CSTPP2 and C11orf49/CSTPP1 knock-out cells, we did not observe nuclear lobulation in TBC1D19 or TTLL1 null cells. Further, LRRC49/CSTPP2 and C11orf49/CSTPP1 knock-outs exhibit persistent ciliation. We therefore deprived *TBC1D19^-/-^* cells of serum and visualized multiple ciliary markers to examine primary cilium assembly. Detection of all these markers indicated that TBC1D19 loss abrogates ciliation, and cilia that assembled were short and stunted (Figs. 4a and S2e). We detected branch-point glutamylation of α-tubulin, needed for the subsequent addition of poly-glutamylate chains, using an antibody, GT335. Ablation of TBC1D19 drastically reduced (by 3-fold) the number of GT335-positive cilia, as compared to the control (Fig. 4a), although total levels of this modified tubulin were not affected (Fig. 3c). Furthermore, the percentage of cells with acetylated tubulin (ac-tubulin)-positive cilia was drastically reduced in the TBC1D19 knockout (Fig. 4a), again consistent with loss of ciliation. Detection of a third well-established ciliary marker, Arl13b, revealed a more dramatic reduction in ciliation (∼12-fold) and altered localization (Fig. 4a). Since total Arl13b levels were not affected (Fig. S4d), our results suggest an intrinsic defect in Arl13b ciliary localization. Loss of primary cilia could not be ascribed to defects in cell cycle progression or an inability to enter a quiescent state (Fig. S2d). We expressed GFP-TBC1D19 in knock-out cells and rescued cilium assembly, again confirming that ciliation defects stemmed from loss of TBC1D19 directly (Fig. S2f). Strikingly, none of these ciliary markers was altered in TTLL1 or C11orf49/CSTPP1 knock-out cells wherein cilia are efficiently assembled (Fig. S2g; (Wang, Paudyal et al. 2022), attesting to unique roles for each of these proteins in holo-TPGC function.

**Figure 4.**
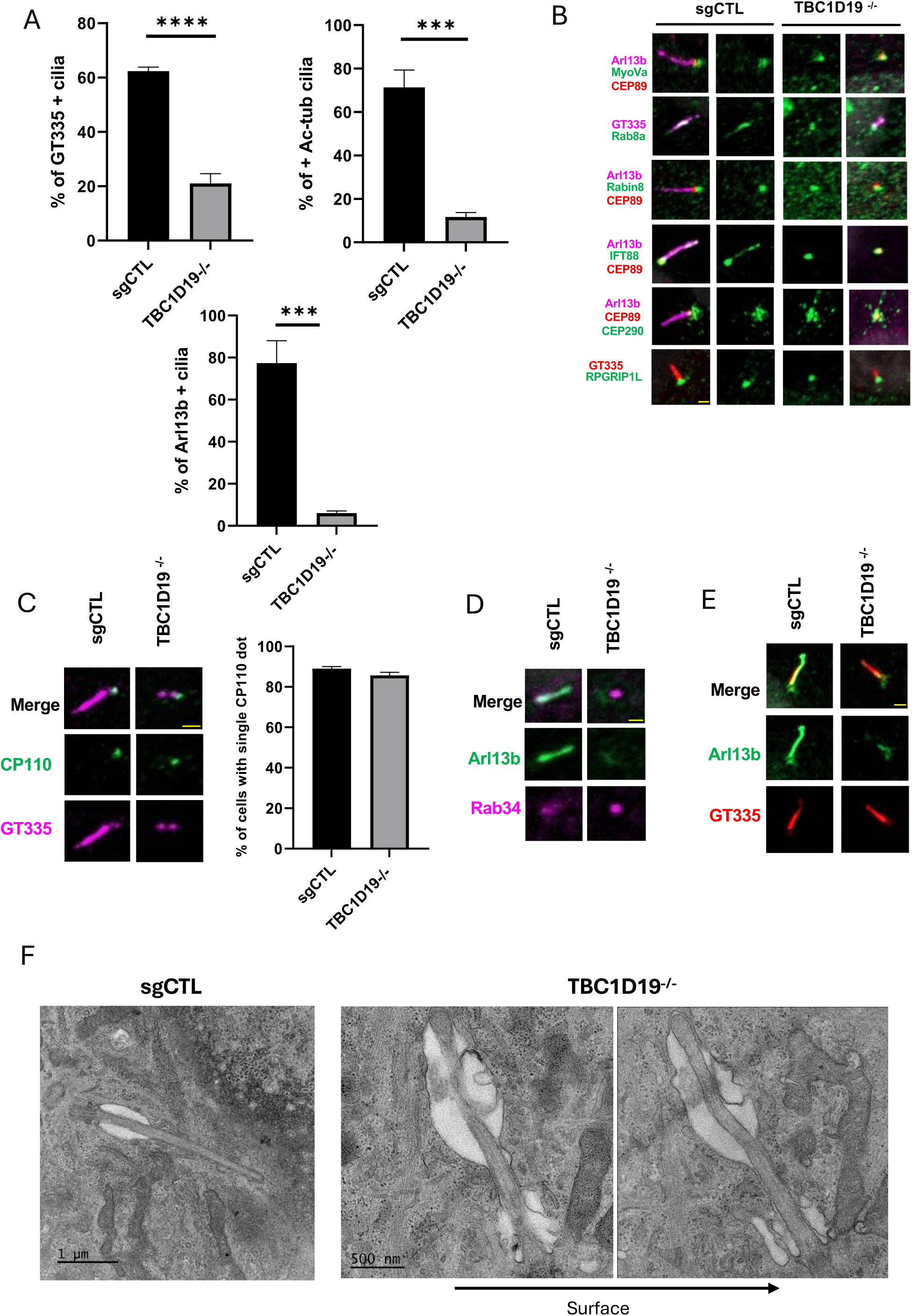
TBC1D19 regulates primary cilium assembly. **A**) sgRNA control and TBC1D19 knockout cells were serum starved for 24 hr and immuno-stained with GT335, anti-acetylated tubulin and anti-Arl13b antibodies. ≥90 cells per sample were analyzed in three independent experiments. Error bars, S.D. ***p ≤ 0.001, **** p ≤ 0.0001. **B**) sgRNA control and TBC1D19 knockout cells were serum starved for 24 hr, and immuno-stained with antibodies against GT335, Arl13b, Myo-Va, Cep89, Rab8a, Rabin8, IFT88, Cep290 and RPGRIP1L. Scale bar, 1µm. **C**) sgRNA control and TBC1D19 knockout cells were serum starved as in panels A and B and visualized by staining with GT335 and antibodies against CP110. Scale bar, 1µm. N ≥ 90 cells per sample were analyzed in three independent experiments. Error bars, S.D. **D**) sgRNA control and TBC1D19 knockout cells were serum starved for 24 hr, and immunostained with antibodies against Rab34 and Arl13b. Scale bar, 1µm. **E**) sgRNA control and TBC1D19 knockout cells were serum starved as in panels A-C and immunostained with antibodies against GT335 and Arl13b. Scale bar, 1µm. **F**) Primary cilia in sgRNA control and TBC1D19 knockout cells were examined by transmission electron microscopy (TEM) after 24 hr of serum starvation. Images of TBC1D19 knockout cilia from two consecutive sections are shown.

Our data suggest that TBC1D19 is essential for assembly of primary cilia, and those cells able to bypass this requirement assemble aberrant or unstable cilia, consistent with the loss of tubulin acetylation and glutamylation, modifications known to stabilize microtubules (Figs. 4 and S2g). Our results also indicate that TBC1D19, TTLL1, and C11orf49/CSTPP1 play surprisingly distinct roles: although each protein is stably associated with TPGC, the loss of each has a dramatically different impact on ciliation.

### TBC1D19 null cells fail to assemble structurally normal cilia

In an effort to pinpoint the mechanistic basis for defective cilium assembly in *TBC1D19*^-/-^ cells, we examined a cohort of proteins associated with each step in the process (Fig. 4b). Since few cilia assembled in this knock-out, and Arl13b was visualized at the base of cilia rather than axonemes (Fig. 4e), our data suggested that ciliation arrests at an early stage. In one of the earliest steps of ciliogenesis, the mother centriole/basal body recruits a ciliary vesicle to the distal appendages (Sanchez and Dynlacht 2016). We found that sub-distal and distal appendage, as well as transition zone proteins that we examined were properly recruited. Likewise, proteins necessary for assembly of the ciliary vesicle, including Rab8, Rabin 8, and MyoVa, were properly recruited to basal bodies (Fig 4b). Further, another signature event, CP110 destruction at mother centrioles, also occurred normally (Spektor, Tsang et al. 2007) (Fig. 4c). We also investigated Rab34, a protein that localizes to the ciliary vesicle and regulates early events in ciliation (Xu, Liu et al. 2018, Oguchi, Okuyama et al. 2020, Ganga, Kennedy et al. 2021, Stuck, Chong et al. 2021) (Fig. 4d). Interestingly, we observed that Rab34 aberrantly accumulated at basal bodies and failed to re-distribute from this location during cilium assembly, unlike control cells, in which the protein disappears as ciliation progresses (Ganga, Kennedy et al. 2021, Stuck, Chong et al. 2021); Fig. S3a), a result consistent with our conclusion that *TBC1D19* null cells fail during the early steps of cilium assembly.

To further investigate possible structural changes in *TBC1D19*^-/-^ cilia, we performed transmission electron microscopy (TEM) on knock-out cells after serum deprivation. Considerably fewer cilia were observed in *TBC1D19*^-/-^ cells as compared to control (sgCTL) cells, whereas control and *C11orf49/CSTPP1*^-/-^ cells assembled mature primary cilia, as expected (Figs. 4f and S3b). We found that *TBC1D19*^-/-^ cilia exhibited several notable structural abnormalities. First, knock-out cells frequently assembled short, abnormally formed nascent cilia with aberrantly shaped ciliary vesicles or more mature, fully formed cilia with extensive vesiculation or membrane blebbing (Figs. 4f and S2e). Upon examination of contiguous sections, we found that portions of *TBC1D19*^-/-^ cilia frequently “disappeared” in one section and reappeared in an adjacent one, in contrast with control cilia that were generally visible in a single plane of focus (Fig. 4f). We conclude that TBC1D19 knock-out cilia exhibited significant curvature. Most notably, *TBC1D19*^-/-^ cilia also had membrane abnormalities that were not observed in control cilia, namely, whereas control cilia were enveloped with a uniform, smooth membrane with a symmetrical ciliary pocket, *TBC1D19*^-/-^ cilia frequently exhibited enlarged, asymmetric ciliary vesicles, pockets, and membranes. The diameters of *TBC1D19*^-/-^ ciliary axonemes were highly variable, and the distal ends were either abnormally bulbous or significantly tapered (Fig. S3c and Video1). The defects at distal ends were reminiscent of cilia derived from cells of ciliopathy patients with *INPP5E* mutations (Bielas, Silhavy et al. 2009, Jacoby, Cox et al. 2009), and *TBC1D19*^-/-^ ciliary membranes were associated with vesicular elements at the proximal end and distal tip. Our results are consistent with the conclusion that TBC1D19 knock-out cells arrest before the growth of a ciliary axoneme, and the population of cells able to bypass this step assemble morphologically abnormal cilia with severe membrane defects.

### TPGC subunits fine-tune microtubule poly-glutamylation and acetylation

Previous work has led to seemingly conflicting conclusions regarding the role of tubulin glutamylation in ciliogenesis and cilium maintenance. Indeed, two tubulin de-glutamylating enzymes, CCP1 (also named AGTPBP1) and the related protein, CCP5/AGBL5, were shown to function either as positive or negative regulators of cilium assembly, respectively (Kim, Lee et al. 2010, Failler, Giro-Perafita et al. 2021). Further, forced removal of glutamylation from microtubules specifically within the axoneme (through expression of the de-glutamylase, CCP5) severely compromised elongation of immature cilia and delayed anterograde, but not retrograde, IFT (Hong, Wang et al. 2018). Improper levels of glutamylation, provoked by loss of either of CCP1 or CCP5, led to destabilization of the distal portion of the axoneme (Mercey, Gadadhar et al. 2024). These observations suggest that tubulin poly-glutamylation is subject to exquisite fine-tuning, and imbalances can result in diverse ciliation defects. We therefore assessed the ability of TPGC subunits to promote tubulin modifications within the centrosome, basal body, and axoneme in cells before and after induction of ciliation. Using our panel of knock-out cell lines, we examined the impact of ablating four different TPGC subunits (TBC1D19, TTLL1, C11orf49/CSTPP1, and LRRC49/CSTPP2). Consistent with previous results (Wang, Paudyal et al. 2022), ablation of C11orf49/CSTPP1 or LRRC49/CSTPP2 triggered a modest decrease in poly-glutamylation at centrioles and ciliary axonemes (Fig. 5a and b). In striking contrast, TBC1D19 and TTLL1 knock-out cells exhibited complete loss of axonemal and centriolar poly-glutamylation (Fig. 5a and b). These results suggest that TPGC is the key centriolar and ciliary poly-glutamylase, and TBC1D19 is critical for assembly and activation of this complex.

Microtubule modifications are thought to be coordinated through a “tubulin code,” although much remains unknown about crosstalk between modifications (Magiera, Singh et al. 2018). We therefore examined other tubulin modifications, including tubulin de-tyrosination. We found that cellular tubulin de-tyrosination levels were significantly reduced in both *TBC1D19^-/-^* and *TTLL1^-/-^* but not *C11orf49*/*CSTPP1^-/-^* cells (Fig. S3d). These findings suggest that residual levels of poly-glutamylation in the latter knock-out may be sufficient to trigger de-tyrosination, whereas a more substantial reduction in poly-glutamylation in *TBC1D19^-/-^* and *TTLL1^-/-^* cells is required to reveal de-tyrosination defects. Biochemical studies have suggested that microtubule de-tyrosination is stimulated by poly-glutamylation *in vitro* (Ebberink, Fernandes et al. 2023), and our findings confirm this link in living cells. Interestingly, we also found that the percentage of cilia (deduced by scoring for IFT88-positivity) marked by acetylated tubulin (ac-tubulin) was significantly reduced in TBC1D19 null cells but not in cells depleted of TTLL1 or C11orf49 (Fig 5c; Wang et al., 2022). In an effort to link axonemal acetylation with ciliary phenotypes, we treated cells with tubacin to inhibit the tubulin de-acetylase, HDAC6, but this treatment did not rescue axonemal acetylation or ciliation (Fig. S3e and f), suggesting that this defect did not stem from aberrant removal of the modification but rather the inability to instigate acetylation in knock-out cells in the first place. Earlier studies have demonstrated a relationship between acetylated-tubulin and microtubule detyrosination (Martinez-Hernandez, Parato et al. 2022). It is possible that the reduction in each of these modifications is linked to the loss of glutamylation, although future studies will be required to further investigate underlying mechanisms.

We used three approaches in an effort to rescue glutamylation in the TBC1D19 null cells. First, since axonemal glutamylation and de-glutamylation are in equilibrium at steady-state, we asked whether tubulin modifications and ciliation can be restored in KO cells after depletion of CCP1, a tubulin de-glutamylase, in TBC1D19 null cells. Depletion of CCP1 enabled partial restoration of tubulin de-tyrosination and ac-tubulin in TBC1D19 null cells (Fig. S4a), concomitant with increased ciliation (Fig. 5d). Our studies suggest that these three modifications may be jointly regulated to help stabilize microtubules, but cross-talk may involve additional regulatory steps. Second, we artificially targeted TTLL1 to microtubules by fusing the enzyme to the MAP4 microtubule-binding domain in an effort to restore glutamylation in TBC1D19 knockout cells. However, expression of this fusion protein was unable to rescue either cellular glutamylation or ciliation in TBC1D19 null cells (Figs. 5e and S4b). This result suggests that targeting TTLL1 to microtubules is not sufficient to restore glutamylation when TBC1D19 has been ablated and TPGC has been compromised. Third, we expressed other monomeric glutamylases, TTLL5 and TTLL6, in TBC1D19 null cells in an effort to rescue glutamylation. TTLL5 is an initiase, adding the branch-point glutamylation, whereas TTLL6 is an elongase. Surprisingly, although we succeeded in rescuing cellular poly- glutamylation levels in TBC1D19 knock out cells through enforced expression of TTLL6 (Fig. S4c), neither TTLL5 nor TTLL6 expression could rescue ciliation in TBC1D19 null cells (Fig. 5f). This result suggests that TPGC-mediated glutamylation is critical for assembly of primary cilia.

**Figure 5.**
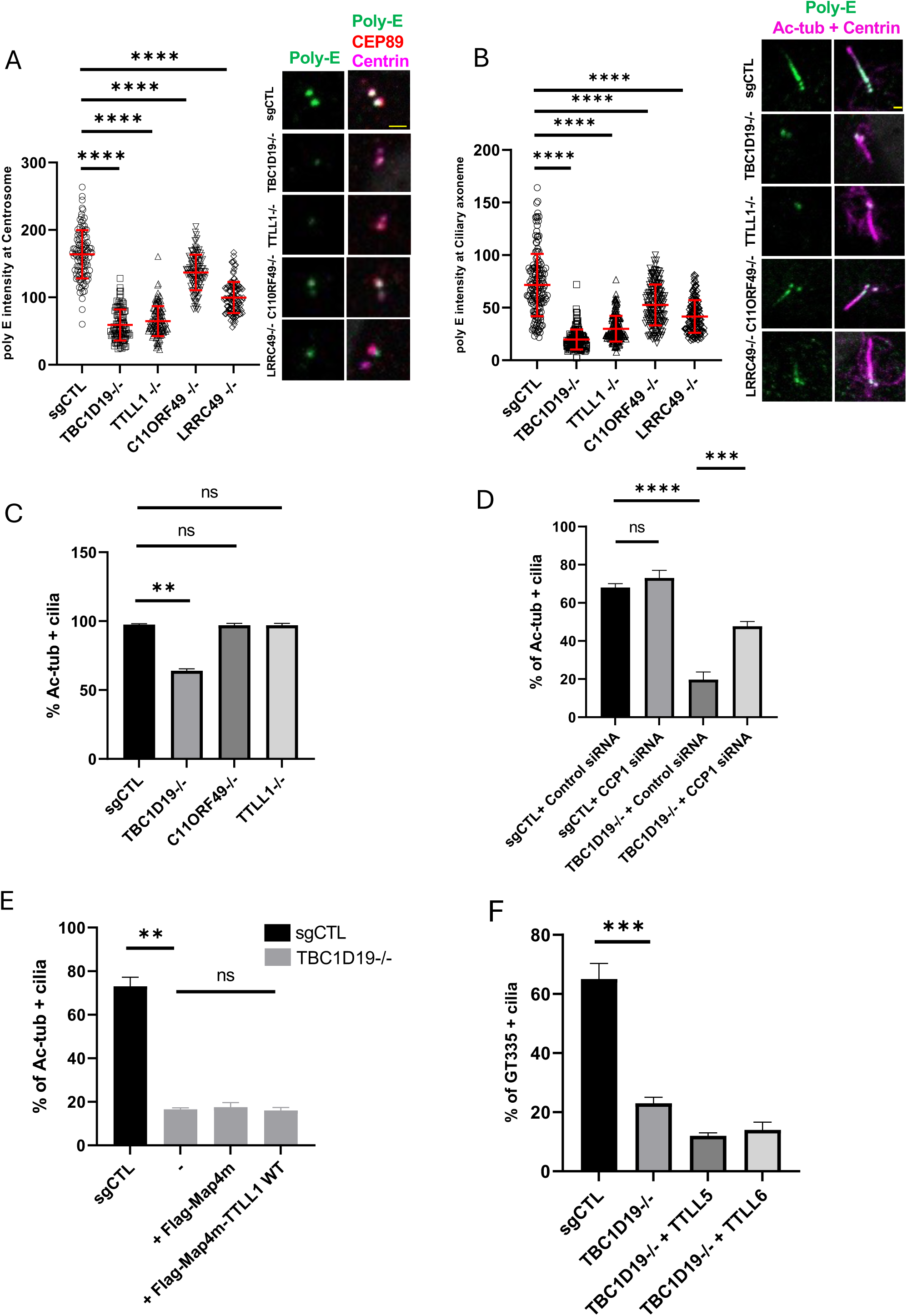
TPGC regulates poly-glutamylation of centriolar and axonemal microtubules. **A**) Centrosomal poly-E intensity in control, *TBC1D19^-/-^*, *TTLL1^-/-^*, *C11ORF49^-/-^*, and *LRRC49^-/-^* cells was measured by staining with antibodies against poly-E, centrin1 and Cep89. G1 cells (two centrin dots) were specifically chosen for quantification. ≥40 cells per sample were counted in two independent experiments. Error bars, S.D. **** p ≤ 0.0001. Representative images are shown for each cell line. Scale bar, 1µm. **B**) Ciliary axonemal poly-E intensity of sgRNA, *TBC1D19^-/-^*, *TTLL1^-/-^*, *C11ORF49^-/-^*, and *LRRC49^-/-^* lines was measured after 24hr serum starvation by staining with antibodies for poly-E, acetylated tubulin, and centrin1. N ≥40 cilia per sample in three independent experiments were counted. Error bars, S.D. **** p ≤ 0.0001. Representative images are shown for each cell line. Scale bar, 1µm. **C**) sgRNA control and TBC1D19 knockout cells were serum starved for 24 hr and immuno-stained with antibodies against IFT88 and acetylated tubulin. N ≥40 IFT88-positive cilia per sample in two independent experiments were counted. Error bars, S.D. **p ≤ 0.01. ns, not significant. **D**) sgRNA control and TBC1D19 knockout cells were treated with control or CCP1 siRNA for 48 hr, serum starved for 24 hr, and immuno-stained with acetylated tubulin antibody. N ≥ 100 cells per sample were counted in three independent experiments. Error bars, S.D. ***p ≤ 0.001, **** p ≤ 0.0001. ns, not significant. **E**) TBC1D19 knockout cells were infected with Flag-map4m or Flag-map4m-TTLL1 lentivirus for 72hr, serum starved for 24 hr, and immunostained with acetylated tubulin antibody. ≥ 90 cells per sample were counted in two independent experiments. Error bars, S.D. **p ≤ 0.01. ns, not significant. **F**) TBC1D19 knockout cells were infected with TTLL5-EYFP or TTLL6-EYFP lentivirus for 72hr, serum starved for 24 hr, and immuno-stained with GT335. N ≥ 90 cells per sample in three independent experiments were counted. Error bars, S.D. ***p ≤ 0.001.

Collectively, these experiments make several important points. First, we conclude that the TPG complex uniquely regulates ciliation via MT poly-glutamylation and that TTLL family members are unable to compensate for loss of TPGC function. Second, since recruitment of TTLL1 to MT is not, by itself, sufficient to restore either glutamylation or ciliogenesis, TPGC components play additional roles beyond MT recruitment. The latter finding comports well with our structural modeling and subsequent experimental data (see below).

### TBC1D19 is essential for INPP5E ciliary entry and cilium stability

Our EM analysis revealed prominent ciliary defects in *TBC1D19^-/-^*cells, most notably, severe ciliary membrane perturbations that could be associated with diminished stability, and we sought a mechanistic explanation for these observations. Phosphoinositides have been shown to play a key role in cilium assembly, stability, and function, and INPP5E, a phosphatidylinositol poly-phosphate 5-phosphatase, is a principal mediator of phosphatidyl inositol homeostasis within the cilium, maintaining low levels of inositol phosphates, PI(4,5)P2 and PI(3,4,5)P3, while enhancing levels of PI(4)P (Bielas, Silhavy et al. 2009, Jacoby, Cox et al. 2009, Garcia-Gonzalo, Phua et al. 2015, Plotnikova, Seo et al. 2015, Phua, Chiba et al. 2019). Loss of INPP5E was shown to affect cilium stability but not its assembly (Bielas, Silhavy et al. 2009, Jacoby, Cox et al. 2009). In other studies, however, INPP5E was shown to be required for axoneme assembly, and displacement of the enzyme during growth induction was associated with cilium decapitation (Phua, Chiba et al. 2019, Sharif, Gerstner et al. 2021). Interestingly, INPP5E ciliary localization was abolished in ciliated TBC1D19 knock-out cells (Fig. 6a), although cellular INPP5E expression levels were not altered (Fig. S4d). We also examined ciliary INPP5E levels in TTLL1, LRRC49, and C11ORF49 knockout cells, and they were unaltered (Figs. 6a and S4e). Since INPP5E plays a demonstrable role in cilium stability, we compared axonemal stability in quiescent control and knock-out cells after short-term serum stimulation. We found that primary cilia in TBC1D19 knock-out cells were substantially less stable, and within 30 minutes of serum addition, ∼40% of cilia had disassembled, as compared to <10% of wild-type cilia (Fig. 6b and c). In contrast, cilium disassembly was induced in *TTLL1-/-* cells with kinetics indistinguishable from controls (Fig. 6d). These findings suggest that cilia inefficiently assemble in TBC1D19 knockout cells and that they are intrinsically unstable during growth induction, but loss of axonemal poly-glutamylation *per se* cannot explain this de-stabilization. Instead, we propose that cilia are de-stabilized in TBC1D19 knockout cells as a result of INPP5E depletion from the organelle.

**Figure 6.**
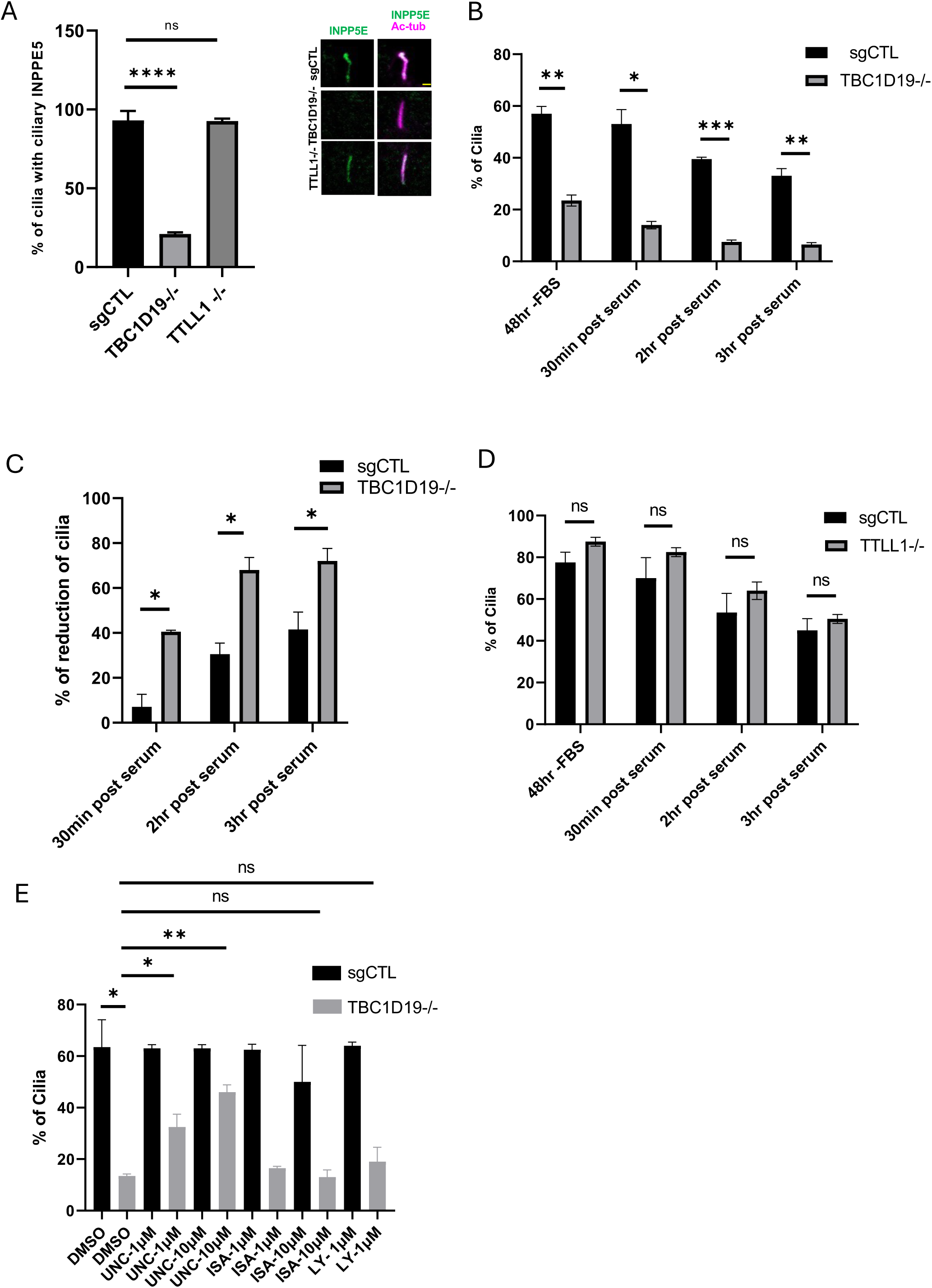
TBC1D19 regulates cilium stability. **A**) sgCTL control, *TBC1D19^-/-^*, and *TTLL1^-/-^* cells were serum starved for 24 hr, and immuno-stained with antibodies against acetylated tubulin and INPP5E. ≥ 80 cilia were counted per sample in three independent experiments. Error bars, S.D. **** p ≤ 0.0001. Representative images were shown for each cell line. Scale bar, 1µm. ns, not significant. **B**) sgRNA control and TBC1D19 knockout cells were serum starved for 48 hr, re-fed with serum-containing medium, fixed after 30 min, 2 hr, or 3 hr, and immuno-stained with GT335 antibody. ≥ 70 cells were counted per sample in two independent experiments. Error bars, S.D. *p ≤ 0.05, **p ≤ 0.01. **C**) Ciliary resorption rate (or percentage reduction in ciliation) after serum addition was calculated based on data in panel B. Error bars, S.D. *p ≤ 0.05. **D**) sgRNA control and TTLL1 knockout cells were treated and stained as in panel B. N ≥ 100 cells were counted per sample in two independent experiments. Error bars, S.D. ns, not significant. **E**) sgRNA control and TBC1D19 knockout cells were serum starved for 24 hr, simultaneously treated with DMSO, 1μM LY-294002 (LY), 1μM or 10 μM UNC3230 (UNC), or 1μM or 10 μM ISA-2011B (ISA), as indicated, and immuno-stained with GT335. ≥ 70 cells were counted per sample in two independent experiments. Error bars, S.D. *p ≤ 0.05, **p ≤ 0.01. ns, not significant.

### Regulation of phosphatidyl inositol homeostasis is a rate-limiting function for TPGC

Our data suggested that loss of INPP5E could explain the ciliary defects observed in TBC1D19 knockout cells. To test this hypothesis, we treated cells with inhibitors of kinases that convert phosphatidyl inositol, PI(4)P, to PI(4,5)P2 and PI(3,4,5)P3, namely, PIP5K1C (phosphatidylinositol-4-phosphate 5-kinase γ) and PI3K, respectively (Vlahos, Matter et al. 1994, Wright, Simpson et al. 2015). Strikingly, treatment of cells with an inhibitor of PIP5K1C (UNC3230), but not PI3K (LY294002), markedly increased ciliation in TBC1D19 knockout cells (Fig. 6e). On the other hand, treatment of cells with ISA-2011-B (Semenas, Hedblom et al. 2014), an inhibitor of the related kinase, PIP5K1A (phosphatidylinositol-4-phosphate 5-kinase α), had no impact (Fig. 6e). These findings are consistent with the observation that PIP5K1C, but not PIP5K1A, localizes to basal bodies (Xu, Zhang et al. 2016) and suggest that aberrant targeting of INPP5E can, in part, explain the loss of ciliation in TBC1D19 knockout cells.

### Loss of TBC1D19 perturbs localization of Arl13b, IFT-A, and ciliary membrane proteins

IFT-B (IFT88) localized normally in TBC1D19 knock-out cells able to ciliate (Fig. 7a). However, the striking exclusion of Arl13b from the sub-population of TBC1D19 knock-out cells able to form cilia prompted us to examine entry of IFT-A (IFT140), since earlier studies have shown the defective ciliary localization of IFT-A in Arl13b knockout cells (Fujisawa, Qiu et al. 2021). We found a significant percentage of TBC1D19 null cilia harbored axonemal IFT140 accumulation compared to other knockouts (Fig. 7b). Moreover, ciliary INPP5E is required for trafficking of SMO and GPR161 to cilia, enabling regulation of the Hedgehog (Hh) pathway (Dyson, Conduit et al. 2017). We examined the transport of a cohort of ciliary membrane proteins, including Smoothened (SMO), and the IFT-A-associated protein, Tulp3, since each of these proteins is able to dock to PI(4,5)P2, and each was shown to be impacted by loss of INPP5E (Garcia-Gonzalo, Phua et al. 2015). We showed that transport of Smo was compromised in *TBC1D19^-/-^* cells able to ciliate after Hh agonist treatment (Fig.7c and Video_1). Interestingly, we found that *TTLL1^-/-^* cells also failed to accumulate Smo, suggesting that glutamylation might play a role in Smo transport into cilia. We also found a significant percentage of cilia with abnormal Tulp3 accumulation in *TBC1D19^-/-^*cells, although in this case, TTLL1 knockout cells did not differ from controls, suggesting that the mechanism for Tulp3 localization to cilia diverges from that of Smo (Fig. 7d). Given the membrane-associated perturbations observed in TBC1D19 null cells (Fig. 7c and d), we examined the localization of another ciliary receptor, MCHR1, in TBC1D19 knockout cells and found that transport of this protein was also significantly reduced (Fig. 7e), suggesting that ciliary membrane protein transport may be generally compromised in the absence of TBC1D19. Gating of proteins by the transition zone was shown to regulate entry of both Arl13b and INPP5E into the cilium (Garcia-Gonzalo, Phua et al. 2015, Yinsheng, Miyoshi et al. 2022). However, we did not detect aberrant localization of the transition zone protein, RPGRIP1L, in the TBC1D19 knock-out (Fig. 4b). Ciliary targeting of INPP5E also requires Arl13b (Humbert, Weihbrecht et al. 2012), and therefore, Arl13b trafficking defects might account for the depletion of INPP5E from cilia in *TBC1D19^-/-^* cells. We posited that, if true, TBC1D19 might be involved in transporting Arl13b into cilia, and we tested this hypothesis by expressing both proteins in 293T cells. Interestingly, Arl13b co-immunoprecipitated with GFP-TBC1D19, suggesting TBC1D19 could help transport Arl13b to cilia (Fig. S4f). Taken together, our data suggest that TBC1D19 is required for Arl13b recruitment, which promotes INPP5E localization essential for stabilization and maintenance of primary cilia. Our findings suggest a model in which TPGC recruits Arl13b to basal bodies, instigating tubulin poly-glutamylation and other tubulin modifications, while facilitating entry of INPP5E into cilia, further promoting cilium stability and proper membrane morphology.

**Figure 7.**
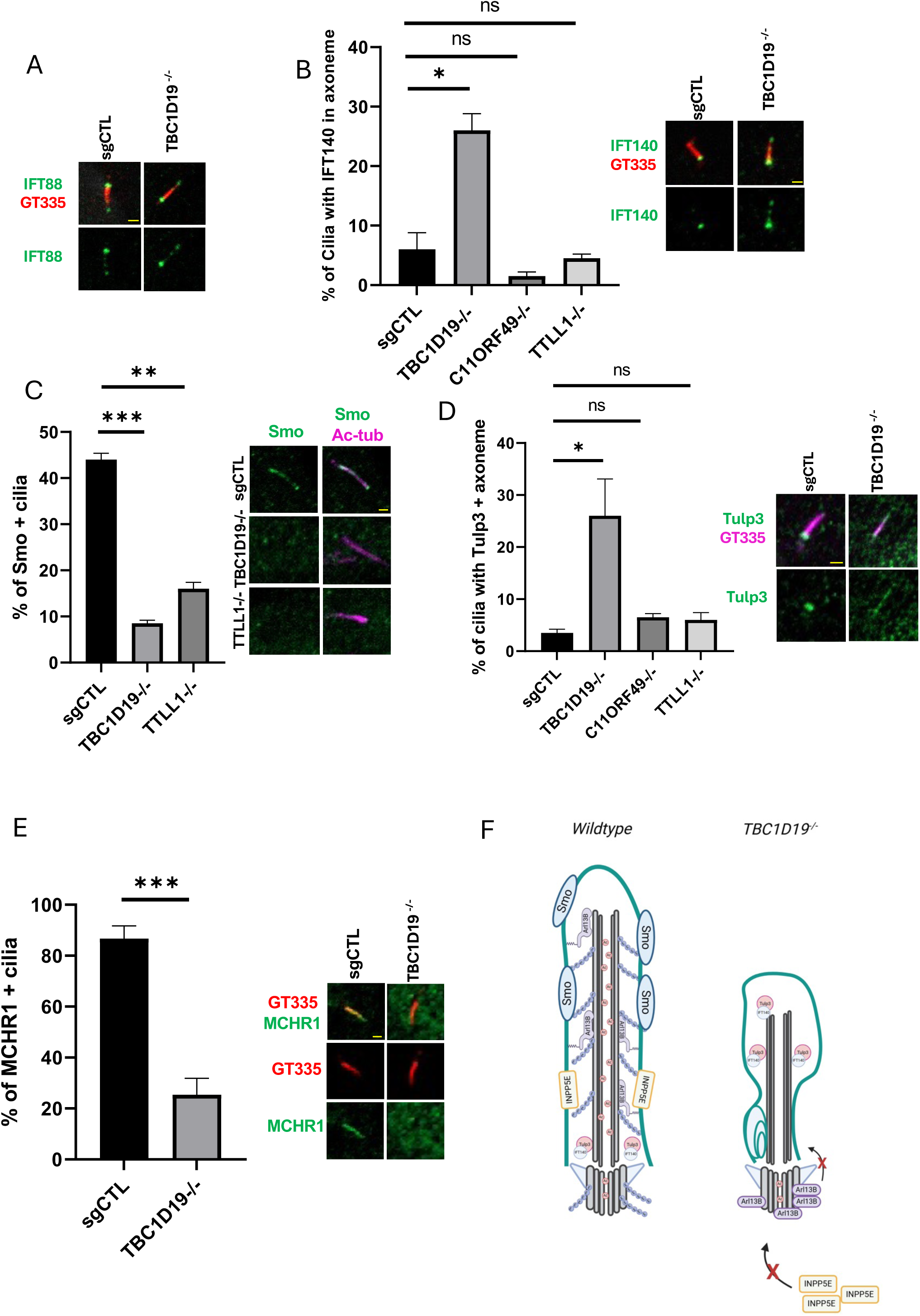
TBC1D19 loss compromises transport of IFT-A and ciliary membrane proteins. **A**) sgRNA control and TBC1D19 knockout cells were serum starved for 24 hr, and immunostained with antibodies against GT335 and IFT88. Scale bar, 1µm. **B**) sgRNA control and TBC1D19 knockout cells, serum starved for 24 hr, and immunostained with antibodies against GT335 and IFT140. Scale bar, 1µm. The percentage of cilia with axonemal IFT140 signal was quantified. N ≥ 45 cilia were counted per sample in two independent experiments. Error bars, S.D. *p ≤ 0.05. Scale bar, 1µm. ns, not significant. **C**) sgRNA control, TBC1D19 knockout and TTLL1 knockout cells were serum starved for 24 hr, simultaneously treated with SAG (200 nm), immunostained with acetylated tubulin and Smo antibodies. Scale bar, 1µm. We quantified the percentage of cilia with axonemal Smo signal. Error bars, S.D. **p ≤ 0.01, ***p ≤ 0.001. N ≥ 70 cilia were counted per sample in two independent experiments. **D**) sgRNA control and TBC1D19 knockout cells were serum starved for 24 hr and immuno-stained with GT335 and antibodies against Tulp3. Scale bar, 1µm. The percentage of Tulp3+ cilia was quantified. Error bars, S.D. *p ≤ 0.05. N ≥ 40 cilia were counted per sample in two independent experiments. ns, not significant. **E**) sgRNA control and TBC1D19 knockout cells stably expressing EGFP-MCHR1 were serum starved for 24 hr and visualized by staining with GT335 and anti-GFP antibodies. Scale bar, 1µm. N ≥ 40 cilia were counted per sample in three independent experiments. Error bars, S.D. ***p ≤ 0.00. **F)** Model showing regulation TBC1D19 in cilium structure, membrane protein transport and ciliary axonemal posttranslational modifications.

Despite its discovery two decades ago (Janke, Rogowski et al. 2005), a complete understanding of the activity and biological function of TPGC has remained elusive. In addition, although we recently identified a sixth component of this complex, C11orf49/CSTPP1 (Wang, Paudyal et al. 2022), we were nevertheless unable to reconstitute robust poly-glutamylase activity with all components, prompting us to identify two novel subunits, TBC1D19 and KIAA1841/SANBR. Here, using gene editing and reconstitution of the enzyme with all eight components, coupled with recently developed protein structure modeling approaches, we have revealed novel insights into the function of this enzyme.

Although our functional studies of KIAA1841 are ongoing, our data suggest a model wherein TBC1D19 and KIAA1841/SANBR are required for the assembly and activation of TPGC (Figs.1 and 2). Once assembled, TPGC initiates steps essential for Arl13b transport, microtubule poly-glutamylation, and INPP5E transport. Our data indicate that TPGC is likely to be required at two stages, first, to promote cilium assembly and secondly, to stabilize the structure once formed (Figs. 4 and 6). Thus, surprisingly, TPGC has been tasked with functions that extend beyond poly-glutamylation (Fig. 7f). Cells that are able to bypass the requirement for INPP5E localization show severe defects in the transport of ciliary membrane proteins and membrane defects, depletion of multiple tubulin modifications (glutamylation, acetylation, and de-tyrosination), triggering organelle instability. Control of phosphatidyl inositol homeostasis and INPP5E localization may represent unanticipated, rate-limiting functions for TBC1D19/TPGC suggestive of a multi-faceted role for TPGC in regulating steps beyond control of tubulin poly-glutamylation. Although other poly-glutamylases, including TTLL5 and TTLL6, among others, are likely to modify axonemal microtubules, our data conclusively show that TPGC is the primary poly-glutamylase regulating centrosomal and ciliary microtubule modifications and, as a consequence, it is likely to be indispensable for organelle assembly and maintenance in some physiological contexts.

We also identified several unanticipated defects in TBC1D19 knock-out cells. In addition to defects in poly-glutamylation, we found that other tubulin modifications, including acetylation and de-tyrosination, were also compromised as a result of TBC1D19 loss. These observations are reminiscent of defects associated with patient mutations in two genes*, ARMC9* and *TOGARAM1*, linked to the neurodevelopmental ciliopathy, Joubert Syndrome, namely, shortened cilia and decreased axonemal acetylation and poly-glutamylation without disruption of transition zone function (Latour, Van De Weghe et al. 2020). Furthermore, mutations in INPP5E have also been linked to Joubert Syndrome (Bielas, Silhavy et al. 2009, Jacoby, Cox et al. 2009). Thus, it will be interesting to explore links between the novel TPGC subunits we have identified and human disease using patient ciliopathy databases.

There are several limitations associated with our studies. For example, we are unable to assess whether the TPGC binds MTs as a monomer or multimer, and while we are able to assemble a tubulin dimer with the holo-TPGC complex, we have not modeled holo-TPGC with multiple protomers or proto-filaments. Further, given that TPGC is an elongase, verification of our models will require structures for co-crystals of TTLL enzymes and the carboxy-terminal tail of α-tubulin modified with the branch-point glutamate, which do not currently exist. We note that *TPGS1* null cells lack α-tubulin glutamylation, although others have shown that β-tubulin glutamylation may also be affected in mouse tissues lacking TTLL1 (Janke, Rogowski et al. 2005, Ikegami, Heier et al. 2007, Ikegami, Sato et al. 2010, Wloga, Joachimiak et al. 2017). It will be interesting to mutate residues within TTLL1 and TPGC based on AlphaFold3 and test how α- and β-tubulin glutamylation is affected in complexes reconstituted with mutant enzymes.

**The authors declare no conflict of interest.**

## Materials and Methods

### Cell culture and generation of knock-out cells

hTERT-RPE-1 and HEK 293T cells were obtained from ATCC. Cells were grown in DMEM or DMEM/F12 medium supplemented with 10% Fetal bovine serum (FBS). Cells were grown in DMEM/F12 without FBS medium for 24 hr or 48 hr for cilia assembly, as indicated. To construct knock-out (KO) cells, RPE-1 cells were infected with Flag-Cas9 and sgRNA expressing lentivirus and propagated for 10 days. Next, cells were seeded in 96-well dishes for single cell isolation, and colonies were analyzed for genome editing. sgRNAs included sgCTL (5’-GAGACGTCTAGCACGTCTCT-3’), sgTBC1D19 (5’-GAGTATCCCACTGGCACGAAA-3’), and sgTTLL1-1 (5’-CCTTCCGGGTCAGTTCATAG-3′). C11ORF49 and LRRC49 KO cells have been described previously (Wang, Paudyal et al. 2022).

### Transfection and lentiviral infection

Polyethylenimine (PEI) was used for transfection of HEK 293T cells at a DNA:PEI ratio of 1:3. HEK 293T cells were incubated for 48 hours prior to collection for protein extraction and immunoprecipitation. Lentivirus was generated in HEK293T cells by co-transfection of the transfer plasmid, Δ8.2 envelope, and VSV-G packaging plasmid using PEI. Lentiviral supernatant was collected after 48-72 hours transfection. RPE1 cells were infected with virus supernatant and 8 μg/ml Polybrene at a 1:1 ratio. Infected cells underwent selection using Geneticin (G418) 500 μg/ml to 1 mg/ml. Lipofectamine RNAiMax (Invitrogen) was used for siRNA transfection of RPE1 cells.

### Plasmids, siRNA, and chemicals

To generate EGFP-tagged TBCD19, the TBC1D19 cDNA (Genescript) was inserted into the pCDH-CMV-Neo vector. pCDH-CMV-FLAG-MAP4m and pCDH-CMV-TTLL1-FLAG-MAP4m were previously described (Wang, Paudyal et al. 2022). To generate EGFP-tagged KIAA1841, KIAA1841 cDNA (Rikon-IRAK167M17) was inserted into the pCDH-CMV-Neo vector. Flag-TTLL1, Flag-TPGS2, Flag-TPGS1, Flag-C11ORF49, Flag-LRRC49, Flag-NICN1 were previously described (Wang, Paudyal et al. 2022). pCDH-TTLL5-YFP and pCDH-TTLL6-YFP (He, Ma et al. 2018) were obtained from J. Hu (Mayo Clinic, Rochester, USA). EGFP-SMOm2 (Wang, Failler et al. 2018). Human centrin 2 was cloned to pCDH-CMV-Neo vector to make pCDH-tagRFP-Centrin-2 plasmid. Flag-C11ORF49 (Wang, Paudyal et al. 2022). To make pLvx-mCherry-Arl13b construct, Arl13b cloned to pLvx-Flag-IRES vector. pWPXLd/LAPC/PURO/ MCHR1 vector obtained from Peter Jackson (Stanford University). siRNAs were purchased from Dharmacon with the following sequences or catalogue numbers: non-specific control (5-AATTCTCCGAACGTGTCACGT-3), CCP1 siRNA -SMARTpool M-014059-00-0005. UNC3230 (T15597) and ISA-2011B (T23498) were purchased from TargetMol. We obtained LY-294002 hydrochloride (L9908) from Sigma and Geneticin G418 from Fisher Scientific (AC329400010). We obtained SAG (HY-12848) and Tubacin (HY-13428) from MedChemExpress.

### Antibodies and immunoprecipitation

We used rabbit anti-Flag (1:2000 for WB and 1:200 for IF, F7425), mouse anti-Flag (1:2000 for WB and 1:500 for IF, F1804), mouse anti-α-tubulin (1:1000 for IF and 1:5000 for WB, T5168), mouse anti-Acetyl-α-Tubulin (1:2000 for IF and 1:1000 for WB, T7451) and rabbit anti-C11ORF49 (1:1000 for WB, HPA040051) from Sigma, goat anti-GFP (1:500 for IF, ab545025,), rabbit anti-LRRC49 (1:1000 for WB, ab189250), rabbit anti-TPGS1 (1:1000 for WB, ab184178) and mouse Anti-detyrosinated α-tubulin antibody [AA12] (1:1500 for IF, ab254154) from Abcam. Mouse anti-α-tubulin (1:10000 for WB 66031-1-Ig), rabbit anti-Arl13b (1:2000 for IF and 1:1000 for WB, 17711-1-AP), rabbit anti-Rab8a (1:50 for IF, 55296-1-AP), rabbit anti-IFT88 (1:500 for IF, 13967-1-AP), rabbit anti-INPP5E (1:1000 for WB and 1:1000 for IF, 17797-1-AP), mouse anti-β-actin (1:10000 for WB, 66009-1-Ig PT), mouse anti-GAPDH (1:10000 for WB, 60004-1-Ig), rabbit ant-CCP1 (1:1000 for WB, 14067-1-AP), Rabbit anti-TBC1D19 (1:1000 for WB, 21085-1-AP) and rabbit anti-Tulp3 (1:1000 for WB, 1:150 for IF, 13637-1-AP), RPGRIP1L (1:150 for IF, 55160-1-AP) were from Proteintech. Rabbit anti-GFP (1:3000 for WB, A-6455) was obtained from Invitrogen. Mouse anti-glutamylated tubulin (GT335) (1:1500 for IF and 1:500 for WB, AG-20B-0020-C100) and rabbit anti-PolyE (1:1000 for IF and 1:500 for WB, A26381411) were obtained from Adipogen.

Mouse anti-PCM1 (1:500 for IF, sc-67204) and mouse anti-Rab34 (1:100 for IF, sc-376710) were obtained from Santa Cruz, Inc. Rabbit anti-detyrosinated-tubulin (1:1000 for WB and 1:1500 for IF, AB3201) and mouse anti-centrin (1:1000 for IF, 04-1624) were obtained from Millipore. Anti-Arl13b (mouse) antibody (1:200 for IF, 180085, Addgene), Rabbit anti-MyoVa (1:500 for IF, NBP1-92156, Novus), and rabbit anti-Cep290 (1: 200 for IF, Bethyl laboratories A301-659A). Other antibodies included rabbit anti-Ofd1 (provided by A. Fry), guinea pig anti-TTLL1 (1:2000 for WB; (Ikegami, Sato et al. 2010), rat anti-CEP89 (Tanos, Yang et al. 2013); gift from Bryan Tsou, MSKCC), Rabin8 (Hattula, Furuhjelm et al. 2002) gift from J. Peranen, University of Helsinki, Finland), CP110 (Chen, Indjeian et al. 2002), rabbit anti-SANBR (1:1000 for WB; (Zheng, Matthews et al. 2021) (gift of B. Vuong, City college of New York), Rabbit anti-TPGS2 (1:4000 for WB: gift from C. Janke, Institute Curie) .

For immunoprecipitations, HEK 293T cells were lysed with ELB buffer (50 mM Hepes, pH 7, 150 mM NaCl, 4 mM EDTA pH 8.0, 0.1% NP40, 10% glycerol, 1 mM DTT, 0.5 mM AEBSF, 2 μg/ml Leupeptin, 2 μg/mL Aprotinin, 10 mM NaF, 50 mM β-glycerophosphate) on ice. Lysates were centrifuged and supernatant collected. Lysate (2-7 mg) was incubated with GFP-Trap Agarose beads (Proteintech). Beads were then washed with ELB buffer and analyzed by immunoblotting.

### Immunofluorescence microscopy

Cells were fixed with cold methanol or 10% formalin solution and permeabilized with 0.03% Triton-X in 1 x PBS. Slides were blocked with 3% BSA in 0.03% Triton-X in PBS before incubation with primary antibodies. Primary antibodies were incubated for 2hr at room temperature. Secondary antibodies used included: AF488 conjugated donkey anti-rabbit IgG (Invitrogen A21206), AF647 conjugated donkey anti-mouse IgG (1:500, Invitrogen A31571), AF488 conjugated donkey anti-goat IgG (1:500, A11055), Cy3-conjugated donkey anti-mouse IgG (1:250, Jackson ImmunoResearch 715-165-150), and Cy3-conjugated donkey anti-rabbit IgG (1:250, Jackson ImmunoResearch 711-165-152). Coverslips were stained with DAPI and mounted onto glass slides with ProLong Gold Antifade reagent (Life Technologies). Images were acquired with a Zeiss LSM 800 confocal microscope (63x, NA 1.4 lens) and analyzed using ImageJ. To calculate intensity of poly-glutamylated tubulin, a region of interest was drawn around centrin-positive foci. This region of interest was then populated onto the poly-E channel, and the mean intensity poly-E signal at the foci was measured. To calculate intensity of poly-glutamylated tubulin (poly-E) within ciliary axonemes, a region of interest was manually drawn through the ciliary axoneme and analyzed using Zeiss Zen lite software.

For live-cell imaging, stable expressing EGFP-SmoM2 + TagRFP-centrin 2 of sgRNA control and TBC1D19-/- knockout cells were seeded in glass bottom dishes and treated with serum free medium for cilium formation. Time Lapse videos were acquired on the Zeiss LSM 800 Confocal microscope. Videos from the Zen software were edited using Microsoft ClipChamp.

### Electron microscopy

Cells were plated on 35mm glass-bottom dishes (P35G-1.5-14-C, MatTek) and fixed with solution containing 2% paraformaldehyde, 2.5% glutaraldehyde, 2mM CaCl_2_ in 0.1 M sodium cacodylate buffer (pH 7.2) at 37°C for 5 minutes, then transferred to fresh fixative and fixed overnight at 4°C. The cells were washed three times, 10 minutes each, post-fixed with 1% osmium tetroxide and 1% potassium ferrocyanide for 1.5 hour on ice, then *en block* stained with 0.25% uranyl acetate aqueous solution overnight at 4°C in the dark. The cells were then washed with ddH_2_O, dehydrated in serially in 30%, 50%, and 70% ethanol on ice for 10 min each, transferring to room temperature, and incubating 10 minutes each with 85%, 90%, and 100% ethanol. Finally, cells were infiltrated and *en face* embedded in Araldilte 502 (Electron Microscopy Sciences). The sample blocks were removed by immersing the dish in liquid nitrogen and trimming under stereoscope. Serial thin (100 nm) sections were cut using a Leica UC6 ultramicrotome, collected on formvar-coated slotted copper grids, and stained with uranyl acetate and lead citrate. Stained grids were examined with a JEOL1400 Flash transmission electron microscope (Japan) and photographed with a Gatan Rio 16 camera (Gatan Inc., Pleasanton, CA).

### Mass spectrometric analysis

Whole-cell extracts were prepared from stable RPE1 cells expressing GFP (control) or GFP- TBC1D19 using ELB buffer as described above, and approximately 50 mg of protein were immuno-purified using GFP-trap agarose chromatography. Mass spectrometric (MS) analysis was performed after tryptic digestion with MS-grade TPCK trypsin (Promega, Madison, WI), and one-third of the supernatant was analyzed per MS run, as previously described (Wang, Paudyal et al. 2022). Experiments were performed in triplicate.

### AlphaFold3 Analysis

#### Inter- and intra-TPGC interaction and 3D structure prediction using AlphaFold3

The amino acid sequences of TPGC members, listed in Table 1, were used to predict the inter- TPGC interactions with AlphaFold3 (Abramson, Adler et al. 2024), and raw data were visualized using PAE viewer (Elfmann and Stulke 2023), PyMOL 3.0.5 and iMovie 10.4.3. The scores of predicted template modeling (pTM) and interface predicted template modeling (ipTM) are indicators of the accuracy of the modeling of intra- and inter-molecular interaction, respectively. A pTM score above 0.5 or higher is considered a good prediction, while the threshold for ipTM is 0.6.

**Table 1.** Uniprot ID of the protein amino acid sequences used for AlphaFold3 simulation.

|  | Uniprot ID |  |
| --- | --- | --- |
|  | Human | Zebrafish |
| LRRC49 | Q8IUZ0 | -- |
| C11ORF49 | Q9H6J7 | Q6DGK9 |
| TBC1D19 | Q8N5T2 | Q7SXU2 |
| TTLL1 | O95922 | A0A8M1P2G8 |
| TPGS1 | Q6ZTW0 | B0S661 |
| TPGS2 | Q68CL5 | Q6GMH5 |
| KIAA1841 (SANBR) | Q6NSI8 | E7EXN1 |
| NICN1 | Q9BSH3 | Q6DEG8 |
| α-tubulin | Q71U36 | Q6NWK7 |
| β-tubulin | Q9H4B7 | Q6P5M9 |

### Statistics

All graphs were plotted using Prism Graph pad software. Statistical significance was calculated using two-tailed unpaired Student’s t-test.

## Supporting information

Video 1a-sgCTL

Video 2_ Holo-TPGC with tubulin dimer

Video 1b-TBC1D19 KO

## Acknowledgments

We thank S. Srivastava for contributions to the characterization of TBC1D19 at the inception of this work. We thank C. Janke for providing reagents and constructive discussions. This work was supported by NIH grant 5R01GM120776 to B.D.D. We thank A. Liang, C. Petzold, and J. Sall at the NYULH Microscopy Lab (partially supported by NYU Cancer Center Support Grant NIH/NCI P30CA016087) for consultation and preparation of EM samples. We thank B. Ueberheide and the NYULH Proteomics Laboratory for assistance with MS analysis. The Proteomics Laboratory is supported in part by NYULH and Perlmutter Cancer Center support grant P30CA016087 from the NCI. We thank P. Jackson, J. Hu, B. Tsou, A. Fry, J. Peranen, and K. Ikegami for antibodies and plasmids. Graphic created (Fig.7f) in BioRender. Collado, L. (2025) https://BioRender.com/o90g778

---

**Fig S1.**
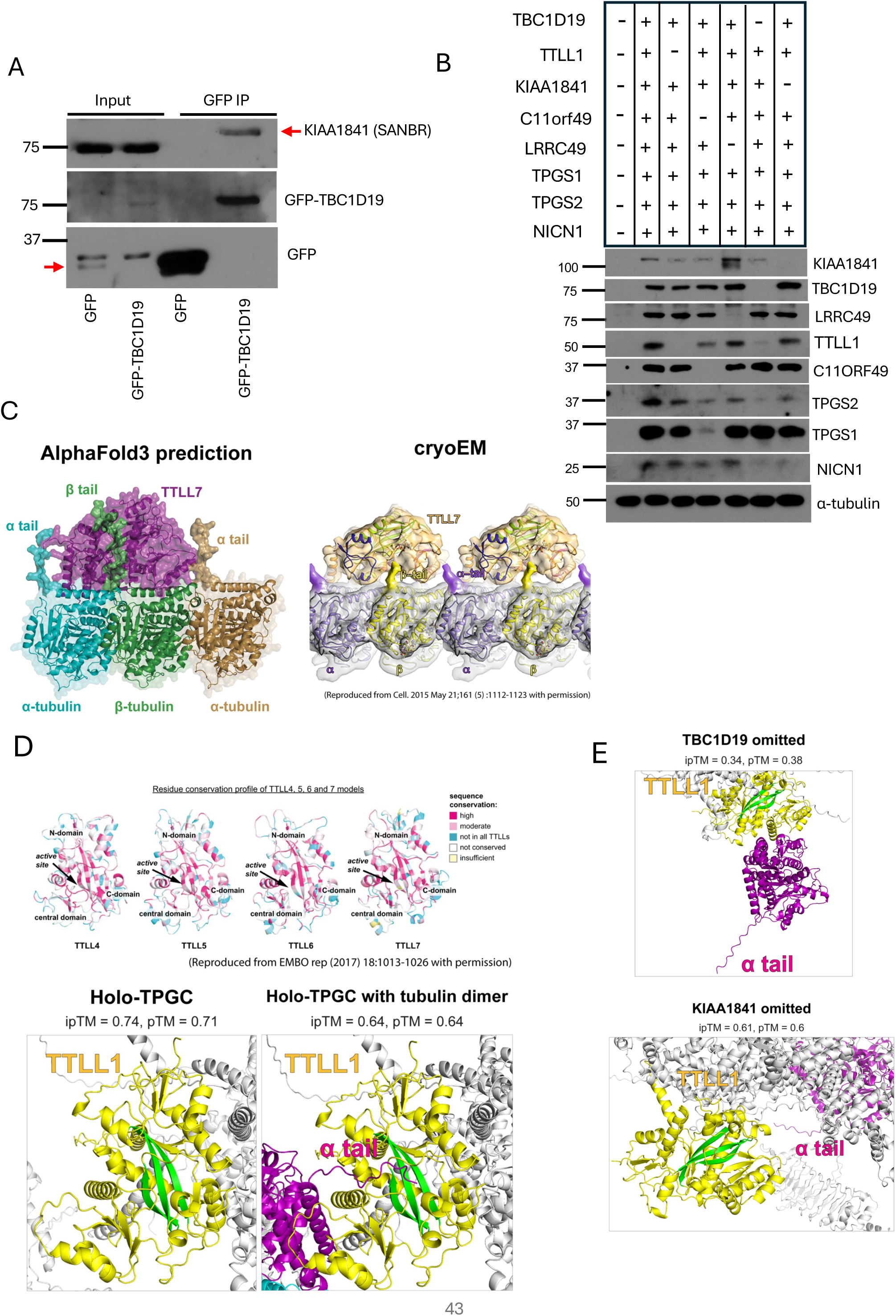
Modeling with AlphaFold3 recapitulates structural data. **A**) RPE-1 cells stably expressing GFP or GFP-TBC1D19 were subjected to immunoprecipitation with GFP-trap beads. Input and immunoprecipitates were immuno- blotted with the indicated antibodies. Note the mobility of KIAA1841/SANBR enriched by GFP-TBC1D19 capture is substantially reduced compared to input, suggesting the presence of post-translational modifications. **B**) Extracts of 293T cells transfected with different combinations of TBC1D19, KIAA1841, TTLL1, NICN1, TPGS2, TPGS1, C11ORF49, and LRRC49, were immunoblotted with antibodies against GFP (to detect TBC1D19 and KIAA1841) and Flag (to detect TTLL1, NICN1, and LRRC49). The remaining proteins were detected with their corresponding antibodies as indicated. **C**) The structure of TTLL7 docked onto tubulin dimer was predicted by AlphaFold3 and compared with published Cryo-EM and crystal structures of TTLL7. The figures were re-published from *Cell* (2015) May 21;161(5):1112-1123 (Graphical abstract) with permission. **D)** We compared AlphaFold3 models of TTLL1 with homology models of TTLL4, 5, 6 and 7 based on crystal structures of tubulin tyrosine ligase (TTL) and glutamylase TTLL7. Structural models of TTLL4, 5, 6 and 7 in the panel were re-published from *EMBO rep* (2017) 18: 1013 – 1026 (Figure 1a) with permission. The β sheets and structural folds of TTLL1 are conserved around the active sites of TTLL4, 5, 6 and 7. **E)** Enlarged view of 3D structures showing the position of α-tubulin tail into holo-TPGC after omission of K1AA1841 or TBC1D19 from the holo-enzyme. Active site β sheets in TTLL1 conserved with TTLL4, 5, 6, and 7 are indicated (green).

**Fig S2.**
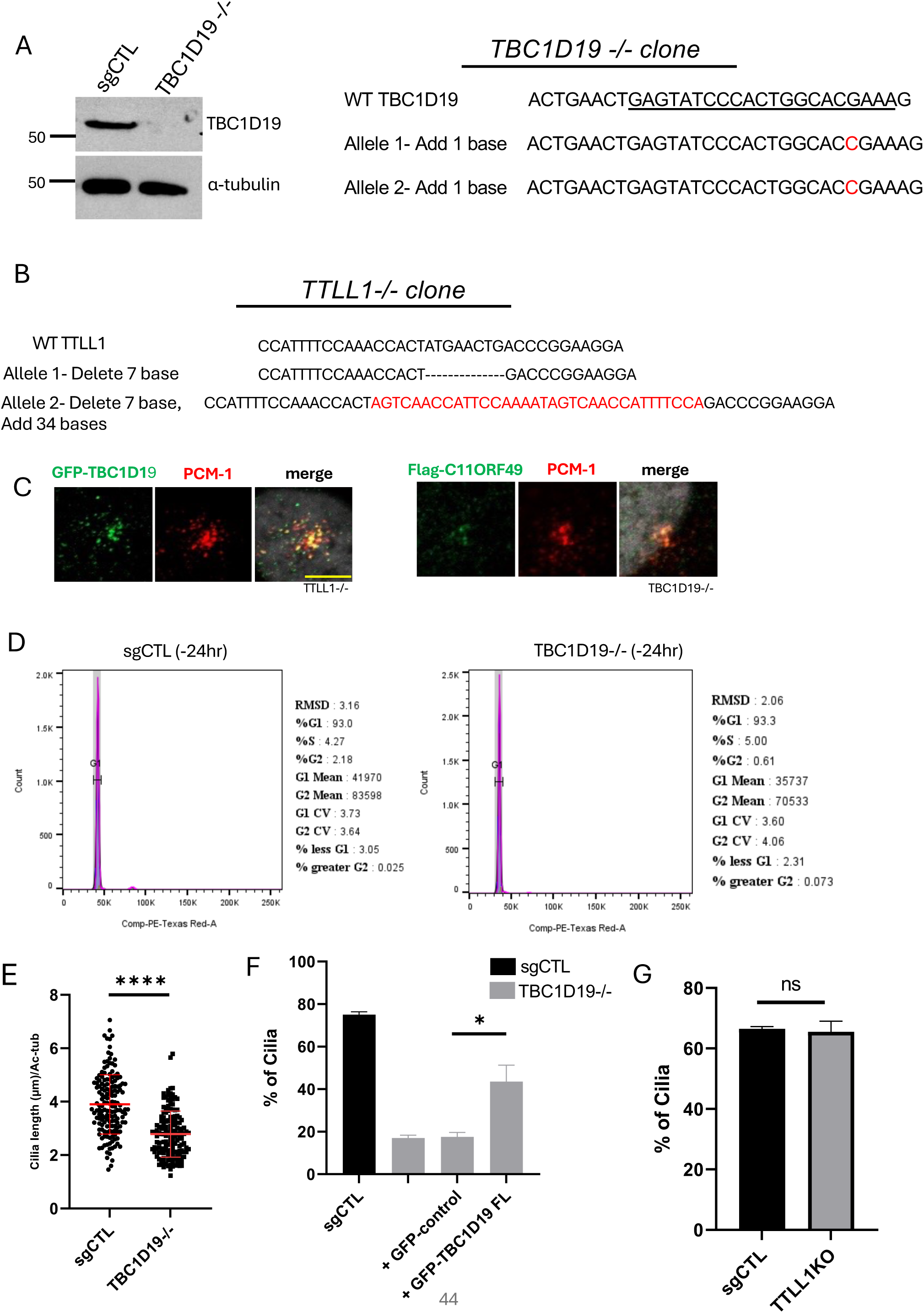
Characterization of *TBC1D19^-/-^* and *TTLL1^-/-^* cells. **A**) sgRNA control and TBC1D19 knockout cells were collected and subjected to immunoblotting with the indicated antibodies. Both alleles of the *TBC1D19* gene targeted by the sgRNA (underlined) were sequenced in the knockout and compared with wild-type genomic region. **B**) The genomic sequence of region targeted by CRISPR in TTLL1 knockout cells is shown and is aligned with wild-type sequence. Deletions and insertions are indicated with dashes and red text, respectively. **C**) TTLL1 knockout cells were infected with lentivirus expressing GFP-TBC1D19 and immuno-stained with antibodies against GFP and PCM-1, and TBC1D19 knockout cells were infected with lentivirus expressing Flag-C11ORF49 and immuno-stained with the same antibodies. Scale bar, 10 µm. **D**) Flow cytometric analysis of sgRNA control and TBCD19 knockout cells stained with propidium iodide after 24 hr of serum starvation indicates G0/G1 arrest. Samples were analyzed in two independent experiments. **E)** sgCTL and TBC1D19 cells were serum starved for 24 hr, and immuno-stained with acetylated tubulin antibody to analyze cilia length. N ≥40 cilia per sample in three independent experiments. Error bars, S.D. **** p ≤ 0.0001. **F**) TBC1D19 knockout cells were infected with lentivirus of GFP-control vector or GFP-TBC1D19 for 72 hr, serum starved for 24 hr, and immuno-stained with acetylated tubulin antibody. N ≥ 90 cells were counted per sample in two independent experiments. Error bars, S.D. ***p ≤ 0.001. **G**) sgRNA control and TTLL1 knockout cells were serum starved for 24 hr and immuno-stained with acetylated tubulin antibody. N ≥ 60 cells were counted per sample in two independent experiments. Error bars, S.D. ns, not significant.

**Fig S3.**
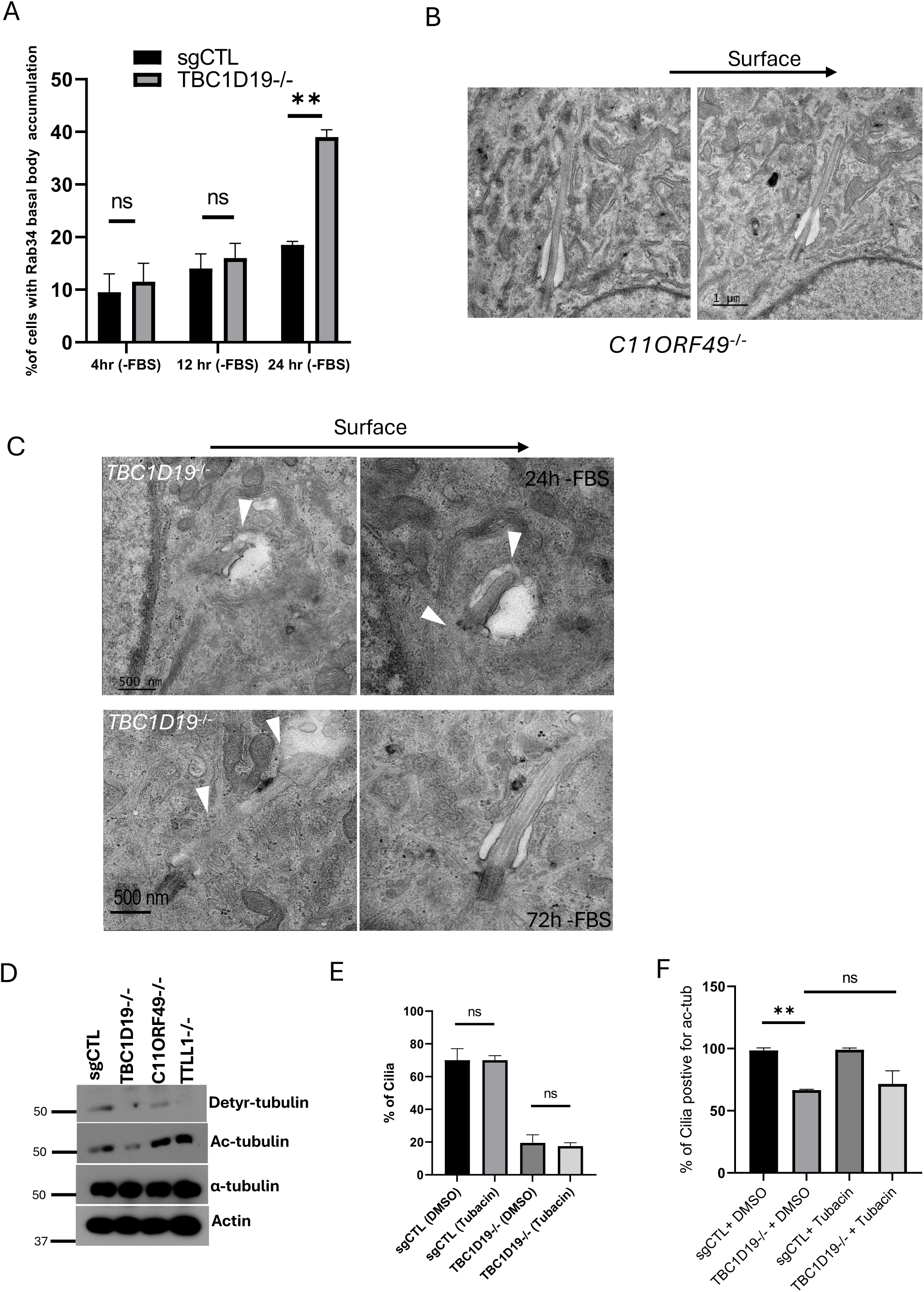
Characterization of ciliary defects in *TBC1D19^-/-^* cells. **A**) Rab34 accumulates at basal bodies in TBC1D19 knockout cells. Cells were stained with antibodies against Rab34 after serum starvation for the indicated period. N ≥ 60 cells were counted per sample in two independent experiments. Error bars, S.D. **p ≤ 0.01. ns, not significant. **B**) Primary cilia in growing C11ORF49 knockout cells were examined by transmission electron microscopy (TEM). Showing adjacent sections of the same cilium. **C**) Primary cilia in TBC1D19 knockout cells were examined by transmission electron microscopy (TEM) after serum starvation (for 24 hr or 72 hr, as indicated). White arrowheads point to bulbous or tapered distal tips, curvature of the axoneme in and out of the plane, or aberrant vesicle formation. **D**) Control (sgCTL), *TBC1D19^-/-^*, *C11ORF49^-/-^*, and *TTLL1^-/-^* cells were serum starved for 24hr and subjected to immunoblotting with the indicated antibodies. **E**) Control (sgCTL) and TBC1D19 knockout cells were serum starved for 24 hr and simultaneously treated with DMSO or Tubacin (2.5 µM). Fixed cells were immuno-stained with antibody against acetylated tubulin. N ≥ 60 cells were counted per sample in two independent experiments. Error bars, S.D. ns, not significant. **F**) sgRNA control and TBC1D19 knockout cells were serum starved for 24hr, simultaneously treated with DMSO or Tubacin (2.5 µM), and immuno-stained with antibodies against acetylated tubulin and IFT88. N ≥ 40 cilia were counted per sample in two independent experiments. Error bars, S.D. **p ≤ 0.01.

**Fig S4.**
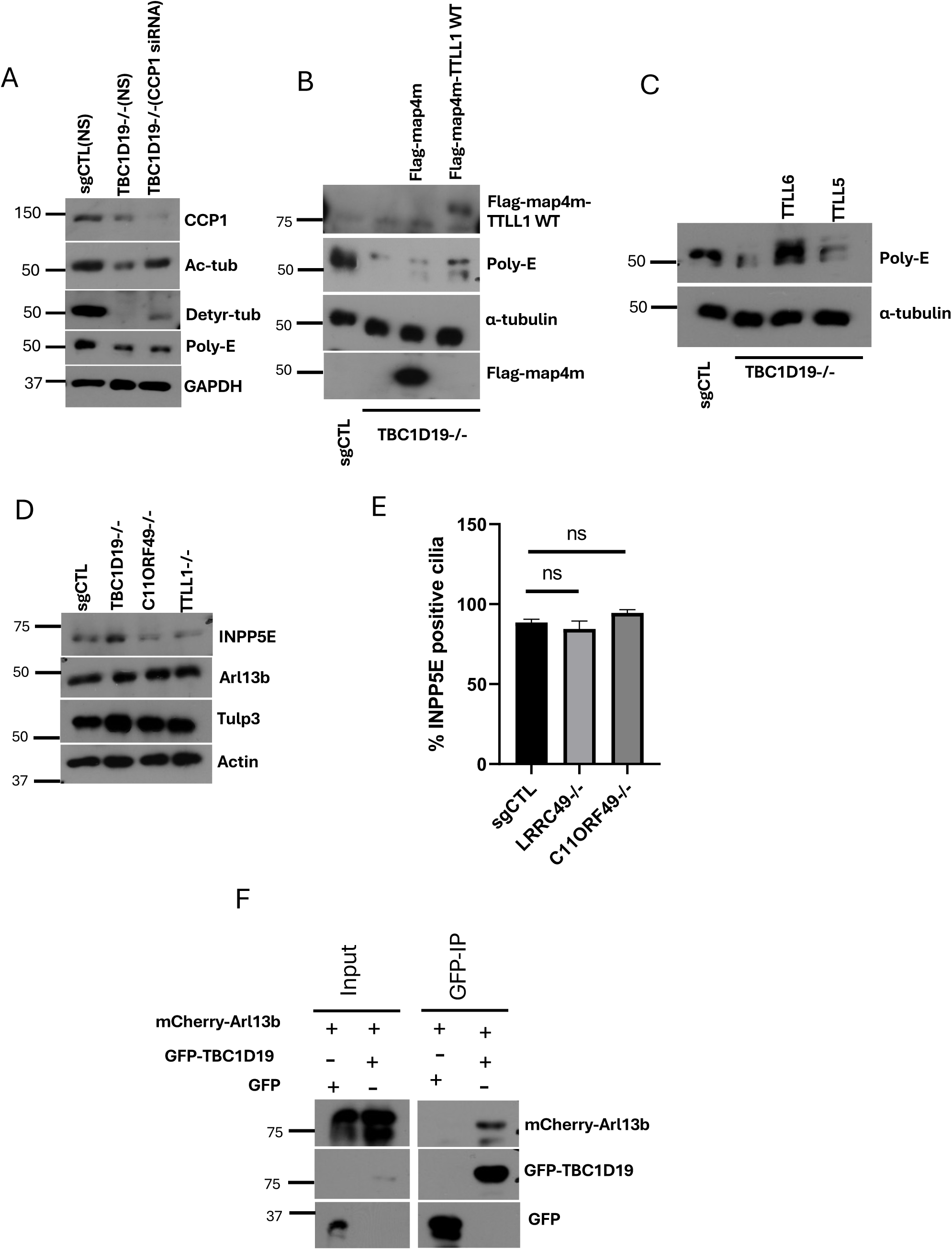
Ablation of TBC1D19, but not other TPGC subunits, triggers loss of Arl13b and INPP5E from cilia. **A**) Control (sgCTL) and TBC1D19 knockout cells were treated with non-targeting control or CCP1 siRNA for 48 hr and serum starved for 24 hr, and extracts were subjected to immunoblotting with the indicated antibodies. **B**) TBC1D19 knockout cells were infected with lentivirus expressing Flag-map4m or Flag-map4m-TTLL1 for 72 hr, serum starved for 24 hr and subjected to immunoblotting with the indicated antibodies. **C**) Control (sgCTL), TBC1D19 knockout cells, and TBC1D19 knockout cells stably expressing TTLL5-EYFP or TTLL6-EYFP were serum starved for 24 hr and subjected to immunoblotting with the indicated antibodies. **D**) Control (sgCTL), *TBC1D19^-/-^*, *C11ORF49^-/-^*, and *TTLL1^-/-^* cells were serum starved for 24 hr and subjected to immunoblotting with the indicated antibodies. **E**) sgCTL, *C11ORF49^-/-^*, and *LRRC49^-/-^* cells were serum starved for 24 hr, immuno-stained with antibodies against acetylated tubulin and INPP5E. N ≥ 50 cilia were counted per sample in two independent experiments. Error bars, S.D. ns, not significant. **F**) 293T cells were co-transfected with GFP or GFP-TBC1D19 with mCherry-Arl13b for 32 hr and serum starved for 16 hr. Lysates were subjected to immunoprecipitation with GFP-trap beads and immuno-blotted with antibodies against GFP and mCherry.

**Video 1a**: Live cell imaging of sgCTL cells stably expressing EGFP-SmoM2 and TagRFP-Centrin-2 after serum starvation.

**Video 1b:** Live cell imaging of *TBC1D19^-/-^*cells stably expressing EGFP-SmoM2 and TagRFP-Centrin-2 after serum starvation.

**Video 2**: 3D structure of human holo-TPGC with tubulin dimer.

## References

Abramson, J., J. Adler, J. Dunger, R. Evans, T. Green, A. Pritzel, O. Ronneberger, L. Willmore, A. J. Ballard, J. Bambrick, S. W. Bodenstein, D. A. Evans, C. C. Hung, M. O’Neill, D. Reiman, K. Tunyasuvunakool, Z. Wu, A. Zemgulyte, E. Arvaniti, C. BeaVe, O. Bertolli, A. Bridgland, A. Cherepanov, M. Congreve, A. I. Cowen-Rivers, A. Cowie, M. Figurnov, F. B. Fuchs, H. Gladman, R. Jain, Y. A. Khan, C. M. R. Low, K. Perlin, A. Potapenko, P. Savy, S. Singh, A. Stecula, A. Thillaisundaram, C. Tong, S. Yakneen, E. D. Zhong, M. Zielinski, A. Zidek, V. Bapst, P. Kohli, M. Jaderberg, D. Hassabis and J. M. Jumper (2024). "Accurate structure prediction of biomolecular interactions with AlphaFold 3." Nature 630(8016): 493–500.

Bielas, S. L., J. L. Silhavy, F. Brancati, M. V. Kisseleva, L. Al-Gazali, L. Sztriha, R. A. Bayoumi, M. S. Zaki, A. Abdel-Aleem, R. O. Rosti, H. Kayserili, D. Swistun, L. C. Scott, E. Bertini, E. Boltshauser, E. Fazzi, L. Travaglini, S. J. Field, S. Gayral, M. Jacoby, S. Schurmans, B. Dallapiccola, P. W. Majerus, E. M. Valente and J. G. Gleeson (2009). "Mutations in INPP5E, encoding inositol polyphosphate-5-phosphatase E, link phosphatidyl inositol signaling to the ciliopathies." Nat Genet 41(9): 1032–1036.

Bonnet, C., D. Boucher, S. Lazereg, B. Pedrotti, K. Islam, P. Denoulet and J. C. Larcher (2001). "Differential binding regulation of microtubule-associated proteins MAP1A, MAP1B, and MAP2 by tubulin polyglutamylation." J Biol Chem 276(16): 12839–12848.

Chen, Z., V. B. Indjeian, M. McManus, L. Wang and B. D. Dynlacht (2002). "CP110, a cell cycle-dependent CDK substrate, regulates centrosome duplication in human cells." Dev Cell 3(3): 339–350.

Dyson, J. M., S. E. Conduit, S. J. Feeney, S. Hakim, T. DiTommaso, A. J. Fulcher, A. Sriratana, G. Ramm, K. A. Horan, R. Gurung, C. Wicking, I. Smyth and C. A. Mitchell (2017). "INPP5E regulates phosphoinositide-dependent cilia transition zone function." J Cell Biol 216(1): 247–263.

Ebberink, E., S. Fernandes, G. Hatzopoulos, N. Agashe, P. H. Chang, N. Guidotti, T. M. Reichart, L. Reymond, M. C. Velluz, F. Schneider, C. Pourroy, C. Janke, P. Gonczy, B. Fierz and C. Aumeier (2023). "Tubulin engineering by semi-synthesis reveals that polyglutamylation directs detyrosination." Nat Chem 15(8): 1179–1187.

Elfmann, C. and J. Stulke (2023). "PAE viewer: a webserver for the interactive visualization of the predicted aligned error for multimer structure predictions and crosslinks." Nucleic Acids Res 51(W1): W404–W410.

Failler, M., A. Giro-Perafita, M. Owa, S. Srivastava, C. Yun, D. J. Kahler, D. Unutmaz, F. J. Esteva, I. Sanchez and B. D. Dynlacht (2021). "Whole-genome screen identifies diverse pathways that negatively regulate ciliogenesis." Mol Biol Cell 32(2): 169–185.

Fujisawa, S., H. Qiu, S. Nozaki, S. Chiba, Y. Katoh and K. Nakayama (2021). "ARL3 and ARL13B GTPases participate in distinct steps of INPP5E targeting to the ciliary membrane." Biol Open 10(9).

Ganga, A. K., M. C. Kennedy, M. E. Oguchi, S. Gray, K. E. Oliver, T. A. Knight, E. M. De La Cruz, Y. Homma, M. Fukuda and D. K. Breslow (2021). "Rab34 GTPase mediates ciliary membrane formation in the intracellular ciliogenesis pathway." Curr Biol 31(13): 2895–2905 e2897.

Garcia-Gonzalo, F. R., S. C. Phua, E. C. Roberson, G. Garcia, 3rd, M. Abedin, S. Schurmans, T. Inoue and J. F. Reiter (2015). "Phosphoinositides Regulate Ciliary Protein Trafficking to Modulate Hedgehog Signaling." Dev Cell **34**(4): 400-409.

Garnham, C. P., A. Vemu, E. M. Wilson-Kubalek, I. Yu, A. Szyk, G. C. Lander, R. A. Milligan and A. Roll-Mecak (2015). "Multivalent Microtubule Recognition by Tubulin Tyrosine Ligase-like Family Glutamylases." Cell 161(5): 1112–1123.

Hattula, K., J. Furuhjelm, A. Arffman and J. Peranen (2002). "A Rab8-specific GDP/GTP exchange factor is involved in actin remodeling and polarized membrane transport." Mol Biol Cell 13(9): 3268–3280.

He, K., X. Ma, T. Xu, Y. Li, A. Hodge, Q. Zhang, J. Torline, Y. Huang, J. Zhao, K. Ling and J. Hu (2018). "Axoneme polyglutamylation regulated by Joubert syndrome protein ARL13B controls ciliary targeting of signaling molecules." Nat Commun 9(1): 3310.

Hong, S. R., C. L. Wang, Y. S. Huang, Y. C. Chang, Y. C. Chang, G. V. Pusapati, C. Y. Lin, N. Hsu, H. C. Cheng, Y. C. Chiang, W. E. Huang, N. C. Shaner, R. Rohatgi, T. Inoue and Y. C. Lin (2018). "Spatiotemporal manipulation of ciliary glutamylation reveals its roles in intraciliary trafficking and Hedgehog signaling." Nat Commun 9(1): 1732.

Humbert, M. C., K. Weihbrecht, C. C. Searby, Y. Li, R. M. Pope, V. C. Sheffield and S. Seo (2012). "ARL13B, PDE6D, and CEP164 form a functional network for INPP5E ciliary targeting." Proc Natl Acad Sci U S A 109(48): 19691–19696.

Huttlin, E. L., R. J. Bruckner, J. Navarrete-Perea, J. R. Cannon, K. Baltier, F. Gebreab, M. P. Gygi, A. Thornock, G. Zarraga, S. Tam, J. Szpyt, B. M. Gassaway, A. Panov, H. Parzen, S. Fu, A. Golbazi, E. Maenpaa, K. Stricker, S. Guha Thakurta, T. Zhang, R. Rad, J. Pan, D. P. Nusinow, J. A. Paulo, D. K. Schweppe, L. P. Vaites, J. W. Harper and S. P. Gygi (2021). "Dual proteome-scale networks reveal cell-specific remodeling of the human interactome." Cell 184(11): 3022–3040 e3028.

Ikegami, K., R. L. Heier, M. Taruishi, H. Takagi, M. Mukai, S. Shimma, S. Taira, K. Hatanaka, N. Morone, I. Yao, P. K. Campbell, S. Yuasa, C. Janke, G. R. Macgregor and M. Setou (2007). "Loss of alpha-tubulin polyglutamylation in ROSA22 mice is associated with abnormal targeting of KIF1A and modulated synaptic function." Proc Natl Acad Sci U S A 104(9): 3213–3218.

Ikegami, K., S. Sato, K. Nakamura, L. E. Ostrowski and M. Setou (2010). "Tubulin polyglutamylation is essential for airway ciliary function through the regulation of beating asymmetry." Proc Natl Acad Sci U S A 107(23): 10490–10495.

Jacoby, M., J. J. Cox, S. Gayral, D. J. Hampshire, M. Ayub, M. Blockmans, E. Pernot, M. V. Kisseleva, P. Compere, S. N. Schiffmann, F. Gergely, J. H. Riley, D. Perez-Morga, C. G. Woods and S. Schurmans (2009). "INPP5E mutations cause primary cilium signaling defects, ciliary instability and ciliopathies in human and mouse." Nat Genet 41(9): 1027–1031.

Janke, C. and J. C. Bulinski (2011). "Post-translational regulation of the microtubule cytoskeleton: mechanisms and functions." Nat Rev Mol Cell Biol 12(12): 773–786.

Janke, C., K. Rogowski, D. Wloga, C. Regnard, A. V. Kajava, J. M. Strub, N. Temurak, J. van Dijk, D. Boucher, A. van Dorsselaer, S. Suryavanshi, J. Gaertig and B. Edde (2005). "Tubulin polyglutamylase enzymes are members of the TTL domain protein family." Science 308(5729): 1758–1762.

Ki, S. M., J. H. Kim, S. Y. Won, S. J. Oh, I. Y. Lee, Y. K. Bae, K. W. Chung, B. O. Choi, B. Park, E. J. Choi and J. E. Lee (2020). "CEP41-mediated ciliary tubulin glutamylation drives angiogenesis through AURKA-dependent deciliation." EMBO Rep 21(2): e48290.

Kim, J., J. E. Lee, S. Heynen-Genel, E. Suyama, K. Ono, K. Lee, T. Ideker, P. Aza-Blanc and J. G. Gleeson (2010). "Functional genomic screen for modulators of ciliogenesis and cilium length." Nature 464(7291): 1048–1051.

Kobe, B. and J. Deisenhofer (1994). "The leucine-rich repeat: a versatile binding motif." Trends Biochem Sci 19(10): 415–421.

Lacroix, B., J. van Dijk, N. D. Gold, J. GuizeV, G. Aldrian-Herrada, K. Rogowski, D. W. Gerlich and C. Janke (2010). "Tubulin polyglutamylation stimulates spastin-mediated microtubule severing." J Cell Biol 189(6): 945–954.

Latour, B. L., J. C. Van De Weghe, T. D. Rusterholz, S. J. Letteboer, A. Gomez, R. Shaheen, M. Gesemann, A. Karamzade, M. Asadollahi, M. Barroso-Gil, M. Chitre, M. E. Grout, J. van Reeuwijk, S. E. van Beersum, C. V. Miller, J. C. Dempsey, H. Morsy, G. University of Washington Center for Mendelian, M. J. Bamshad, C. Genomics England Research, D. A. Nickerson, S. C. Neuhauss, K. Boldt, M. Ueffing, M. Keramatipour, J. A. Sayer, F. S. Alkuraya, R. Bachmann-Gagescu, R. Roepman and D. Doherty (2020). "Dysfunction of the ciliary ARMC9/TOGARAM1 protein module causes Joubert syndrome." J Clin Invest **130**(8): 4423-4439.

Lee, G. S., Y. He, E. J. Dougherty, M. Jimenez-Movilla, M. Avella, S. Grullon, D. S. Sharlin, C. Guo, J. A. Blackford, Jr., S. Awasthi, Z. Zhang, S. P. Armstrong, E. C. London, W. Chen, J. Dean and S. S. Simons, Jr. (2013). "Disruption of Ttll5/stamp gene (tubulin tyrosine ligase-like protein 5/SRC-1 and TIF2-associated modulatory protein gene) in male mice causes sperm malformation and infertility." J Biol Chem 288(21): 15167–15180.

Lee, J. E., J. L. Silhavy, M. S. Zaki, J. Schroth, S. L. Bielas, S. E. Marsh, J. Olvera, F. Brancati, M. Iannicelli, K. Ikegami, A. M. Schlossman, B. Merriman, T. AVe-Bitach, C. V. Logan, I. A. Glass, A. Cluckey, C. M. Louie, J. H. Lee, H. R. Raynes, I. Rapin, I. P. Castroviejo, M. Setou, C. Barbot, E. Boltshauser, S. F. Nelson, F. Hildebrandt, C. A. Johnson, D. A. Doherty, E. M. Valente and J. G. Gleeson (2012). "CEP41 is mutated in Joubert syndrome and is required for tubulin glutamylation at the cilium." Nat Genet 44(2): 193–199.

Magiera, M. M., P. Singh, S. Gadadhar and C. Janke (2018). "Tubulin Posttranslational Modifications and Emerging Links to Human Disease." Cell 173(6): 1323–1327.

Martinez-Hernandez, J., J. Parato, A. Sharma, J. M. Soleilhac, X. Qu, E. Tein, A. Sproul, A. Andrieux, Y. Goldberg, M. J. Moutin, F. Bartolini and L. Peris (2022). "Crosstalk between acetylation and the tyrosination/detyrosination cycle of alpha-tubulin in Alzheimer’s disease." Front Cell Dev Biol 10: 926914.

Mercey, O., S. Gadadhar, M. M. Magiera, L. Lebrun, C. Kostic, A. Moulin, Y. Arsenijevic, C. Janke, P. Guichard and V. Hamel (2024). "Glutamylation imbalance impairs the molecular architecture of the photoreceptor cilium." EMBO J 43(24): 6679–6704.

Natarajan, K., S. Gadadhar, J. Souphron, M. M. Magiera and C. Janke (2017). "Molecular interactions between tubulin tails and glutamylases reveal determinants of glutamylation patterns." EMBO Rep 18(6): 1013–1026.

Oguchi, M. E., K. Okuyama, Y. Homma and M. Fukuda (2020). "A comprehensive analysis of Rab GTPases reveals a role for Rab34 in serum starvation-induced primary ciliogenesis." J Biol Chem 295(36): 12674–12685.

Phua, S. C., S. Chiba, M. Suzuki, E. Su, E. C. Roberson, G. V. Pusapati, S. Schurmans, M. Setou, R. Rohatgi, J. F. Reiter, K. Ikegami and T. Inoue (2019). "Dynamic Remodeling of Membrane Composition Drives Cell Cycle through Primary Cilia Excision." Cell 178(1): 261.

Plotnikova, O. V., S. Seo, D. L. Cottle, S. Conduit, S. Hakim, J. M. Dyson, C. A. Mitchell and I. M. Smyth (2015). "INPP5E interacts with AURKA, linking phosphoinositide signaling to primary cilium stability." J Cell Sci 128(2): 364–372.

Sanchez, I. and B. D. Dynlacht (2016). "Cilium assembly and disassembly." Nat Cell Biol 18(7): 711–717.

Semenas, J., A. Hedblom, R. R. Mioakhova, M. Sarwar, R. Larsson, L. Shcherbina, M. E. Johansson, P. Harkonen, O. Sterner and J. L. Persson (2014). "The role of PI3K/AKT-related PIP5K1alpha and the discovery of its selective inhibitor for treatment of advanced prostate cancer." Proc Natl Acad Sci U S A 111(35): E3689–3698.

Sharif, A. S., C. D. Gerstner, M. A. Cady, V. Y. Arshavsky, C. Mitchell, G. Ying, J. M. Frederick and W. Baehr (2021). "Deletion of the phosphatase INPP5E in the murine retina impairs photoreceptor axoneme formation and prevents disc morphogenesis." J Biol Chem 296: 100529.

Sharma, N., J. Bryant, D. Wloga, R. Donaldson, R. C. Davis, M. Jerka-Dziadosz and J. Gaertig (2007). "Katanin regulates dynamics of microtubules and biogenesis of motile cilia." J Cell Biol 178(6): 1065–1079.

Sirajuddin, M., L. M. Rice and R. D. Vale (2014). "Regulation of microtubule motors by tubulin isotypes and post-translational modifications." Nat Cell Biol 16(4): 335–344.

Spektor, A., W. Y. Tsang, D. Khoo and B. D. Dynlacht (2007). "Cep97 and CP110 suppress a cilia assembly program." Cell 130(4): 678–690.

Stuck, M. W., W. M. Chong, J. C. Liao and G. J. Pazour (2021). "Rab34 is necessary for early stages of intracellular ciliogenesis." Curr Biol 31(13): 2887–2894 e2884.

Szczesna, E., E. A. Zehr, S. W. Cummings, A. Szyk, K. K. Mahalingan, Y. Li and A. Roll-Mecak (2022). "Combinatorial and antagonistic effects of tubulin glutamylation and glycylation on katanin microtubule severing." Dev Cell 57(21): 2497–2513 e2496.

Tanos, B. E., H. J. Yang, R. Soni, W. J. Wang, F. P. Macaluso, J. M. Asara and M. F. Tsou (2013). "Centriole distal appendages promote membrane docking, leading to cilia initiation." Genes Dev 27(2): 163–168.

Valenstein, M. L. and A. Roll-Mecak (2016). "Graded Control of Microtubule Severing by Tubulin Glutamylation." Cell 164(5): 911–921.

van Dijk, J., K. Rogowski, J. Miro, B. Lacroix, B. Edde and C. Janke (2007). "A targeted multienzyme mechanism for selective microtubule polyglutamylation." Mol Cell 26(3): 437–448.

Vlahos, C. J., W. F. Matter, K. Y. Hui and R. F. Brown (1994). "A specific inhibitor of phosphatidylinositol 3-kinase, 2-(4-morpholinyl)-8-phenyl-4H-1-benzopyran-4-one (LY294002)." J Biol Chem 269(7): 5241–5248.

Vogel, P., G. Hansen, G. Fontenot and R. Read (2010). "Tubulin tyrosine ligase-like 1 deficiency results in chronic rhinosinusitis and abnormal development of spermatid flagella in mice." Vet Pathol 47(4): 703–712.

Wang, L., M. Failler, W. Fu and B. D. Dynlacht (2018). "A distal centriolar protein network controls organelle maturation and asymmetry." Nat Commun 9(1): 3938.

Wang, L., S. C. Paudyal, Y. Kang, M. Owa, F. X. Liang, A. Spektor, H. Knaut, I. Sanchez and B. D. Dynlacht (2022). "Regulators of tubulin polyglutamylation control nuclear shape and cilium disassembly by balancing microtubule and actin assembly." Cell Res 32(2): 190–209.

Wloga, D., E. Joachimiak, P. Louka and J. Gaertig (2017). "Posttranslational Modifications of Tubulin and Cilia." Cold Spring Harb Perspect Biol 9(6).

Wright, B. D., C. Simpson, M. Stashko, D. Kireev, E. A. Hull-Ryde, M. J. Zylka and W. P. Janzen (2015). "Development of a High-Throughput Screening Assay to Identify Inhibitors of the Lipid Kinase PIP5K1C." J Biomol Screen 20(5): 655–662.

Xu, Q., Y. Zhang, Q. Wei, Y. Huang, J. Hu and K. Ling (2016). "Phosphatidylinositol phosphate kinase PIPKIgamma and phosphatase INPP5E coordinate initiation of ciliogenesis." Nat Commun 7: 10777.

Xu, S., Y. Liu, Q. Meng and B. Wang (2018). "Rab34 small GTPase is required for Hedgehog signaling and an early step of ciliary vesicle formation in mouse." J Cell Sci 131(21).

Yinsheng, Z., K. Miyoshi, Y. Qin, Y. Fujiwara, T. Yoshimura and T. Katayama (2022). "TMEM67 is required for the gating function of the transition zone that controls entry of membrane-associated proteins ARL13B and INPP5E into primary cilia." Biochem Biophys Res Commun 636(Pt 1): 162–169.

Zheng, S., A. J. Matthews, N. Rahman, K. Herrick-Reynolds, E. Sible, J. E. Choi, A. Wishnie, Y. K. Ng, D. Rhodes, S. J. Elledge and B. Q. Vuong (2021). "The uncharacterized SANT and BTB domain-containing protein SANBR inhibits class switch recombination." J Biol Chem 296: 100625.

